# Adaptive changes of cholinergic projections to the nucleus accumbens bidirectionally mediate cocaine reinforcing effects

**DOI:** 10.1101/2025.06.30.662309

**Authors:** LAA Aguiar, L Royon, R Bastos-Gonçalves, TTA Carvalho, E Teixeira, AC Faria, RM Jungmann, D Vilasboas-Campos, AV Domingues, C Soares-Cunha, NAP Vasconcelos, L Pinto, SP Fernandez, J Barik, B Coimbra, AJ Rodrigues

**Affiliations:** Life and Health Sciences Research Institute (ICVS), School of Medicine, University of Minho, Braga, Portugal; ICVS/3B’s–PT Government Associate Laboratory, Braga/Guimarães, Portugal; Institut de Pharmacologie Moléculaire & Cellulaire, CNRS, UMR7275, Valbonne, France; Université Côte d’Azur, Nice, 06560, France; Inserm U1323; Departamento de Engenharia Biomédica, Universidade Federal de Pernambuco, Brazil

**Author notes:** corresponding authors: Requests for further information and resources should be directed to and will be fulfilled by the lead contacts, Bárbara Coimbra and Ana João Rodrigues. contributed equally (LAA Aguiar, L Royon, C Soares-Cunha).

**Keywords:** Cocaine, mesopontine, nucleus accumbens, laterodorsal tegmentum, acetylcholine, cocaine sensitization

## Abstract

The laterodorsal tegmentum (LDT) sends critical inputs to distinct reward circuit regions, including the nucleus accumbens (NAc), but their functional role in addiction-related behaviors remains underexplored. Here, we demonstrate that LDT-NAc cholinergic projections undergo cocaine-induced adaptations and modulate cocaine-related behaviors. Using cell type-specific tracing, we show that LDT neurons preferentially innervate NAc medium spiny neurons and cholinergic interneurons. Large-scale *in vivo* recordings reveal that cocaine pre-exposure induces persistent alterations in both LDT and NAc neuronal dynamics and modifies responses to subsequent cocaine challenge. Remarkably, pre-exposure to cocaine triggers population-specific adaptations in the LDT, selectively enhancing excitability of LDT-NAc-projecting cholinergic neurons while reducing that of non-projecting cholinergic cells. Behaviorally, optogenetic activation of LDT-NAc cholinergic projections enhances cocaine conditioning, whereas their inhibition diminishes cocaine’s reinforcing effects.

Our findings identify LDT-NAc cholinergic inputs as key substrates of cocaine-induced plasticity, and critical mediators of cocaine’s rewarding properties, introducing a novel component to addiction circuitry.

## Introduction

The laterodorsal tegmentum (LDT) is a conserved brainstem nucleus that serves as a critical node in ascending arousal pathways. Located in the dorsal pons, this brain region has been canonically related to arousal, consciousness and REM sleep ^1–3^. Recently, the LDT has attracted more attention due to its involvement in reward and reinforcement ^4–8^. The LDT contains segregated cholinergic, glutamatergic, and GABAergic neurons ^4,9,10^, which densely innervate the ventral tegmental area (VTA), regulating the activity of dopamine ^11–13^, and GABA neurons ^14,15^. By modulating the firing of midbrain dopamine neurons, the LDT consequently influences dopamine release in the nucleus accumbens (NAc)^13,16^, a mechanism thought to contribute to the rewarding effects of natural rewards and substances of abuse. Moreover, LDT activity is essential for VTA dopamine neurons burst firing^13^, which is considered a functionally relevant signal for positive reinforcement^17^. In agreement, optogenetic stimulation of LDT-VTA neurons induces place preference^5,18^ and intra-cranial self-stimulation in rats^7,19^. Beyond this modulatory influence of VTA activity, the LDT also directly innervates the NAc^4,10^, a key region in reward and reinforcement. Importantly, LDT-NAc cholinergic inputs shift preference and enhance motivation to work for food rewards^4^.

Despite these findings, and the extensive research on cocaine’s neural mechanisms, the specific role of the LDT-NAc circuitry in mediating cocaine responses remains unknown. This knowledge gap is particularly surprising given the *ex vivo* evidence revealing profound and persistent neuronal modifications in both the LDT and NAc following cocaine exposure. For example, repeated cocaine exposure enhances excitatory synaptic transmission in the LDT and changes intrinsic membrane plasticity ^20,20,21^. Similarly, cocaine triggers synaptic changes in NAc neurons due, with potentiation of medium spiny neurons expressing the D1 dopamine receptor (D1-MSNs) ^22–24^. However, to date, we lack comprehensive insight into how brief cocaine pre-exposure alters the LDT-NAc circuitry, and the relative contributions of cholinergic, GABAergic, or glutamatergic inputs to cocaine behavioral effects.

In this work, we performed large scale *in vivo* electrophysiological recordings of LDT and NAc neurons in a model of subchronic cocaine sensitization, that consists in exposing *naïve* mice to five days of cocaine. Despite the relatively brief exposure regimen, this model presents cocaine-locomotor sensitization, and recapitulates the emergence of a negative affective state following cocaine exposure, presenting reward deficits, and anhedonia^22,25,26^. Moreover, given the scarse knowledge about the electrophysiological signature of LDT neuronal subtypes, we also performed optotagging to unambiguously identify distinct neuronal populations. We further integrated *in vivo* data with *ex vivo* patch-clamp recordings, and optogenetic manipulations of specific LDT-NAc projecting neurons during cocaine conditioning.

Our data shows that pre-exposure to cocaine triggers changes in specific neuronal populations of the LDT and NAc, profoundly altering the responses to an acute cocaine challenge. Notably, our findings clearly pinpoints to a prominent role of LDT-NAc cholinergic inputs in mediating cocaine’s reinforcing properties.

## Results

### LDT preferentially innervates MSNs and cholinergic interneurons

The LDT sends inputs to the NAc, though it is still unknown to which type of cells it is connected to. Therefore, we used anterograde and retrograde viral tracing strategies to characterize LDT projections to the NAc. First, we injected a cre-dependent anterograde virus in the LDT (AAV5-DiO-ChR2-eYFP) of ChAT-cre, vGluT2-cre and vGAT-cre animals to evaluate the presence of YFP^+^ terminals in the NAc. Our data shows that the NAc contains cholinergic, glutamatergic and GABAergic LDT axonal terminals (Figure 1A).

**Figure 1.**
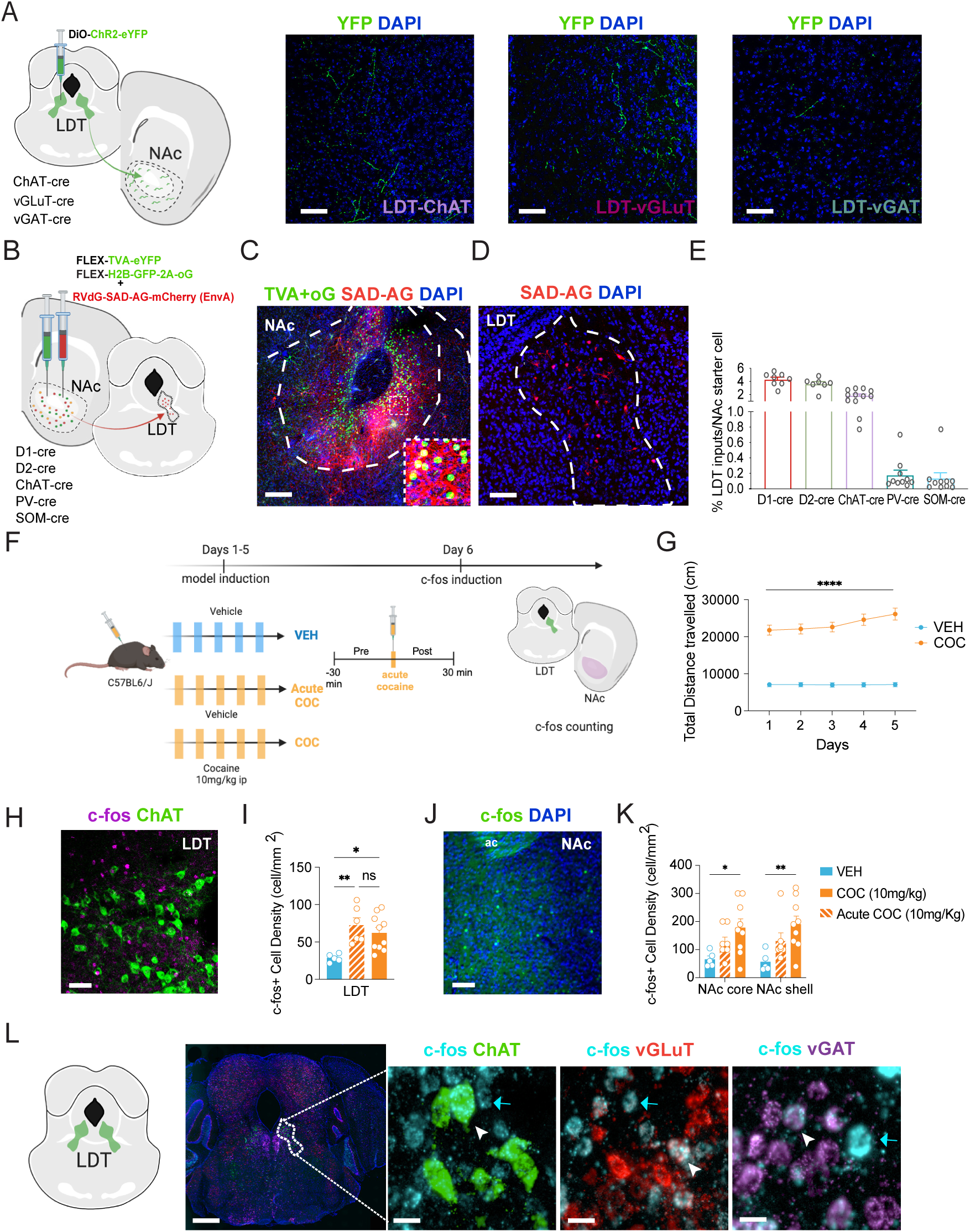
LDT sends cholinergic, glutamatergic and GABAergic inputs to the NAc and is activated by repeated or acute cocaine exposure. **A)** Schematic representation of the anterograde tracing strategy. A cre-dependent ChR2-YFP virus was injected in the LDT of ChAT-cre, vGluT2-cre or vGAT-cre mice, in order to analyze the presence of YFP^+^ terminals in the NAc. In the NAc (right images), we found evidence of cholinergic, glutamatergic and GABAergic LDT terminals. Scale bar, 100µm. B) Scheme of the monosynaptic rabies tracing strategy. We injected in the NAc of D1-cre, D2-cre, ChAT-cre, SOM-cre and PV-cre animals, helper viruses driving the expression of TVA receptor and G protein (with GFP tag) and two weeks later, the G-deleted rabies virus that has a mCherry tag. C) Representative image of a NAc showing co-infected cells labeled in yellow (GFP^+^+mCherry^+^; Scale bar, 250µm; inset showing double positive starter cells). D) LDT cells connected to the NAc cells will be labeled with mCherry; Scale bar, 150µm. E) LDT neurons project to the main five neuronal populations of the NAc, although preferentially innervating D1- and D2-MSNs and cholinergic interneurons (D1: 8 animals; D2: 7 animals; ChAT: 12 animals; PV: 10 animals; SOM: 10 animals). F) Experimental timeline. We measured the density c-Fos^+^ cells in order to evaluate the recruitment of LDT and NAc regions upon repeated or acute cocaine exposure. For this, three experimental groups were created: COC group animals received five daily injections of cocaine (10 mg/kg, i.p.); VEH group received five daily injections of vehicle; Acute COC animals received a single cocaine (10 mg/kg, i.p.) injection. Animals were sacrificed 1h after cocaine or vehicle injection. G) Repeated exposure to cocaine (COC group) triggered a progressively enhanced locomotor response (n=10 animals per group). H) Representative image of the LDT region, showing the characteristic ChAT^+^ cells and c-Fos^+^ cells. Scale bar, 50µm. I) Quantification of c-Fos^+^ cell density in the LDT in VEH, COC and acute COC groups. Acute or repeated cocaine exposure increases the density of c-fos^+^ cells in the LDT (n=5 animals; 8-10 slices). J) Representative image of the NAc, showing c-Fos^+^ cells. K) Quantification of c-fos^+^ cell density in the NAc in VEH, COC and acute COC groups. Repeated cocaine exposure increases the density of c-fos^+^ cells in the NAc (n=5 animals; 8-10 slices). L) RNAScope hybridization showing that cocaine recruits cholinergic, glutamatergic and GABAergic cells, as shown by the co-localization of c-Fos^+^ cells with ChAT^+^, vGLut2^+^ and vGAT^+^, respectively (white arrows; blue arrows mark cells that are c-Fos^+^ only. Scale bar, 50µm. Data represented as mean±SEM. *p < 0.05, **p < 0.01, ***p < 0.001.

Next, to identify to which NAc neurons LDT neurons make monosynaptic contacts, we used the G-deleted rabies tracing approach^27^ to identify targeted NAc neuronal populations. For this, we injected in the NAc of D1-cre, D2-cre, ChAT-cre, SOM-cre and PV-cre animals, helper viruses driving the expression of TVA receptor and G protein (with GFP tag) and two weeks later, the G-deleted rabies virus (mCherry tag) (Fig.1B). NAc co-infected cells will be labeled in yellow (co-expressing GFP and mCherry; Fig. 1C), and monosynaptically connected cells from the LDT (or elsewhere in the brain) will be labeled with mCherry (Fig. 1D). Interestingly, LDT neurons target all of the five neuronal populations analyzed of the NAc, although preferentially innervating D1- and D2-MSN and cholinergic interneurons (CINs) (Fig. 1E, 1way ANOVA, F = 67.89, p < 0.0001).

In sum, the main LDT neuronal populations directly innervate the NAc, and synapse into all subtypes of NAc neurons analyzed, though preferentially with MSNs and CINs.

### Cocaine recruits LDT and NAc cells

Next, we sought to determine whether prior subchronic exposure to cocaine would alter neuronal responses to an acute cocaine challenge. For this, we pre-exposed naïve mice to an i.p. injection of cocaine for 5 days (COC) (and a respective control group receiving 5 days of saline - VEH). An additional group of naïve mice was included to evaluate the effects of an acute cocaine injection (Acute COC). First, we validated the subchronic cocaine sensitization model, by evaluating cocaine-induced locomotor sensitization. As depicted in Figure 1G, in comparison to VEH, COC animals presented increased locomotion in response to cocaine throughout the 5 days of exposure (2way ANOVA; F(4, 68) = 43.92, p<0.0001). After model induction, on the 6^th^ day, all groups were exposed to a single i.p. injection of cocaine. Animals were sacrificed 1-hour post-administration of cocaine, and brains were processed for c-Fos quantification, as an index of neuronal recruitment (Figure 1F). We analysed the number of recruited cells (c-Fos^+^) in the LDT and NAc. COC and acute groups present increased number of c-Fos^+^ cells in the LDT (Fig. 1H-I; LDT: 1way ANOVA, p 0.0089, Bonferroni post-hoc comparison: VEH vs COC p=0.0291; VEH vs Acute COC p=0.0089). No changes were found between acute and COC groups regarding c-Fos expression. In the NAc, cocaine pre-exposure increased the density of c-Fos^+^ cells when compared to VEH animals, in both core and shell subregion (Fig. 1J-K; NAc: 2way ANOVA, p=0.0152; Bonferroni post-hoc comparison: VEH vs COC NAc core: p=0.0292; NAc core: p=0.0075).

Next, we aimed to disentangle which LDT cells were being recruited in response to cocaine, as this nucleus contains cholinergic, glutamatergic, and GABAergic neurons ^4,9,10^. For this, we performed RNAScope for each LDT cell using *ChAT*, *vGlut2* and *VGAT* probes in combination with *c-fos* probe. Importantly, all types of neurons were recruited in the LDT in response to cocaine (Fig. 1L).

Altogether, these results show that repeated and acute cocaine recruits both the LDT and the NAc regions, with differences in the magnitude of the effect.

### Cocaine pre-exposure alters LDT and NAc neuronal activity

To understand the impact of the short regimen of repeated cocaine exposure in the LDT and NAc regions, we performed electrophysiological recordings in COC and VEH anesthetized animals using dense silicon probe recordings in both structures 24h after the last cocaine (or saline) injection (Figure 2A-B; probe placement in Figure S1A). After recording baseline activity (*pre* period) for at least 1h, we administered cocaine 10 mg/kg i.p. and recorded for an additional hour (*post* period). For simplification purposes, we only present 30 min before and after cocaine exposure.

**Figure 2.**
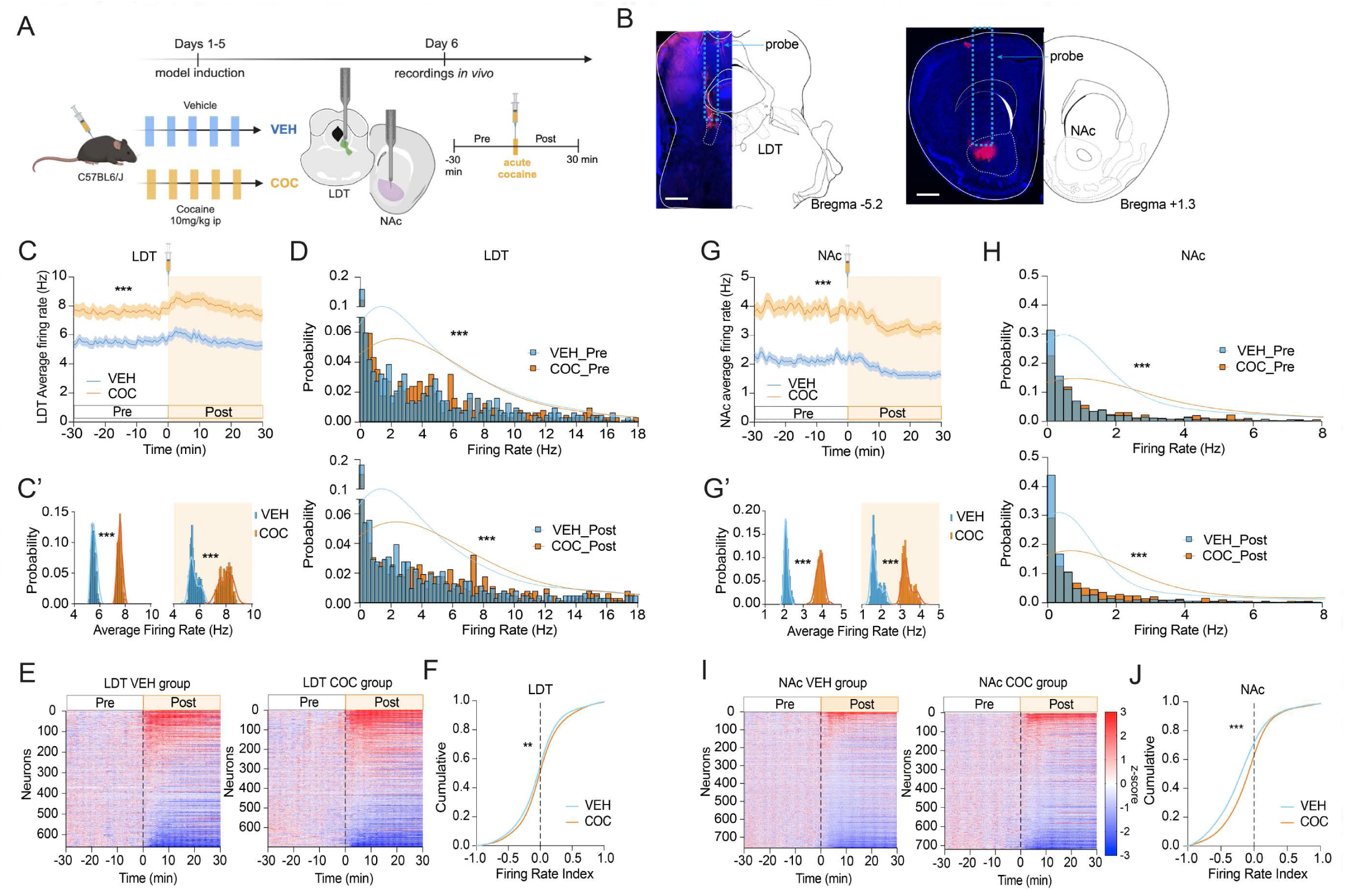
Pre-exposure to cocaine triggers neuroadaptations in the LDT and in the NAc. **A)** Experimental timeline. Animals in the COC group received five daily injections of cocaine (10 mg/kg, i.p.); VEH group received vehicle injections. On the 6th day, animals were subjected to acute electrophysiological recordings in the LDT and NAc using 64 channel silicone probes. After recording baseline period (pre), animals were challenged with an acute injection of cocaine 10 mg/kg, i.p. (post period). **B)** Representative image of a DiI-soaked probe inserted in the LDT and in the NAc (staining red). Scale bar, 500µm. Blue dashed line depicts probe placement. **C)** PSTH depicting the average firing rate of the LDT for the COC (orange, n=706) and VEH (blue, n=651) groups. Cocaine was administered at 0 timepoint. **C’** Distribution of the average firing rate in VEH and COC groups in the pre and post cocaine periods. Pre-exposure to cocaine increases average basal firing rate of the LDT. **D)** Distribution of the individual firing rates in the LDT during the 30-minute window before (top) and after (bottom) acute cocaine injection. COC group presents a significantly different distribution in comparison to VEH group. **E)** Heatmap showing the responses of individual neurons of the LDT for the VEH and COC groups during the 30 minutes before and after acute cocaine injection. The black vertical line represents the time of cocaine administration. Data are z-scored to baseline activity, and organized by response to cocaine. **F)** Distribution of the firing rate index in VEH and COC groups. Firing rate is the difference in activity between post and pre periods for each individual neuron. No significant differences were found. **G)** PSTH depicting the average firing rate of the NAc for the COC (orange, n=699) and VEH (blue, n=660) groups. Cocaine was administered at 0 timepoint. Pre-exposure to cocaine increases NAc average basal firing rate. **H)** Distribution of the individual firing rates in the NAc during the 30-minute window before and after acute cocaine injection. COC group presents a significantly different distribution in comparison to VEH group. **I)** Heatmap showing the responses of NAc individual neurons for the VEH and COC groups during the 30 minutes before and after acute cocaine injection. **J)** Distribution of the firing rate index of the NAc in VEH and COC groups. There was a shift to the right in COC group. PSTH data represented as mean±SEM. *p < 0.05, **p < 0.01, ***p < 0.001.

We recorded hundreds of neurons of each brain region; only single units were included in the analysis (n_LDT-VEH_=651, n_LDT-COC_=699). Interestingly, in basal conditions, the LDT net average firing rate of the COC group was higher than in VEH animals (Figure 2C; Mann-Whitney, p < 0.0001); average baseline firing rate of COC = 7.6 Hz, VEH = 5.5 Hz). When observing the distribution of the average activity in the pre and post cocaine periods, this was significantly different between COC and VEH groups; (Figure 2C’, Pre: Mann-Whitney U=0.0, p < 0.0001; Post: Mann-Whitney, p < 0.0001); as well as the distribution of the firing rate of all recorded cells (Figure 2D, Pre: Mann-Whitney U=215814.5000, p < 0.0001; Post: Mann-Whitney, p < 0.0001). Importantly, these basal differences in activity were still present after 15 days of withdrawal, revealing long-lasting cocaine-induced changes in LDT neuronal dynamics (Figure S2A-B).

Next, we evaluated the effects of an acute cocaine injection. Cocaine increased the average firing rate of the LDT in both VEH and COC animals, with marked magnitude in the first 10 minutes after administration, as it can be observed in the peristimulus time histogram (PSTH) (Figure 2C, VEH: Mann-Whitney U = 902.0, p < 0.0001; COC: Mann-Whitney U = 34.64, p < 0.0001). Analyzing individual units showed that there were cells that presented increased activity in response to cocaine, but there was also a subset of cells that were inhibited in response to cocaine in both groups (Figure 2E; pie charts with % of each type of response in Figure S1B). To further understand the impact of acute cocaine on LDT neuronal responses, we calculated the firing rate index, which is the difference in firing rate between post and pre periods for each individual neuron. Remarkably, acute cocaine effects were more pronounced in COC group, supported by a rightward shift of the firing index curve in relation to VEH group (Figure 2F; Mann-Whitney U=222296.5000, p<0.0001).

Regarding the effects in the NAc, we found that cocaine pre-exposure increased the basal net activity of this region (n_NAc-VEH_=755; n_NAc-COC_=711; Figure 2G-G’). The average activity and the distribution of the firing rate of the NAc in COC group was higher than in VEH group (VEH = 2.2 Hz vs COC = 3.9 Hz; Mann-Whitney U=211756.0000, p=0,0055; Figure 2G’, Pre: Mann-Whitney U = 0.0, p < 0.0001; Post: Mann-Whitney U = 0.0, p < 0.0001).

Conversely to LDT data, acute cocaine injection decreased the average firing rate of the NAc in both groups, with pronounced effects in the first 10 minutes after cocaine administration (Figure 2G; VEH: Mann-Whitney U = 10398.0, p < 0.0001; COC: Mann-Whitney U = 11011.0, p < 0.0001). The analyses of individual units showed that there was also a subset of cells that were excited in response to cocaine in both groups (Fig. 2I; pie charts with % of each type of response in Figure S1C). The firing index was significantly different between COC and VEH groups (Figure 2J; Mann-Whitney U= 214730.00, p<0,0001).

These findings reveal that cocaine pre-exposure triggers long-lasting changes in LDT and NAc neuronal activity and primes these brain regions to respond differently to a subsequent cocaine challenge. Given the direct anatomical and functional connectivities between LDT and NAc, cocaine pre-exposure may fundamentally alter how LDT activity influences NAc neuronal dynamics. Since we recorded both regions simultaneously, we used Reduced Rank Regression (RRR) to identify low-dimensional neural communication patterns between these two brain areas (see methodology for details). We trained a single RRR model on the first 5 min of baseline data, then applied it to successive 5 min windows before and after injection. Prediction quality was evaluated via coefficient of determination (r²) between true and predicted firing-rate indices in each window; allowing to assess how well the baseline-trained model continued predicting NAc activity over time. During pre-injection, both VEH and COC groups maintained high r^2^ values (0.90–0.95), demonstrating stable cross-regional communication structure during baseline conditions (Fig. S2C). Following cocaine injection, both groups show a reduction in r^2^ values, confirming that cocaine substantially disrupts LDT-NAc circuit dynamics. Critically, this reduction was more pronounced in the COC group, indicating that cocaine pre-exposure fundamentally reorganizes the communication between these regions (Fig. S2C; 10 to 15: Mann-Whitney U = 196.0, p = 0.0287; 15 to 20: Mann-Whitney U = 206.0, p = 0.011; 20 to 25: Mann-Whitney U = 210.0, p = 0.007; 25 to 30: Mann-Whitney U = 221.0, p = 0.001).

Collectively, these results demonstrate that subchronic cocaine exposure alters individual regional activity, and significantly alters LDT-NAc functional connectivity.

### Cocaine pre-exposure alters the electrophysiological signature of distinct LDT neuronal clusters

Our previous findings show that pre-exposure to cocaine remodels the LDT-NAc circuitry. Building on these observations, we next sought to delineate the neuronal population(s) mediating these circuit-level adaptations. Albeit there are studies performing *ex vivo* electrophysiological recordings of mesopontine regions (LDT/PPN), *in vivo* recordings in these regions are scarce^28^, and the electrophysiological characteristics of LDT subpopulations are still not defined. Thus, our first goal was to identify the electrophysiological signature of distinct LDT neuronal populations *in vivo*. For this, we used WaveMAP tool^29^, an unsupervised analysis method that clusters cells according to their average extracellular action potential waveforms. We complemented this with CellExplorer tool, that calculates standardized physiological metrics and performs neuron-type classification based on 22 electrophysiological functional features^30^. To complement and validate these methods, we also injected a cre-dependent AVV5-DIO-ChR2-eYFP in either ChAT-cre, VGlut2-cre and vGAT-cre animals, which allows optotagging^31^, in order to unambiguously validate our signature map with genetically-identified cells.

WaveMAP identified six clusters of neurons based on waveform characteristics (we only included cells from VEH animals; Figure S3A). When optotagged cells were added to the UMAP, we observe that cholinergic, glutamatergic and GABAergic cells were distributed throughout more than one cluster, indicating diversity within each neuronal population (Figure S3B-D). Confirming previous findings, using diverse combinations of CellExplorer parameters, we found that GABAergic, glutamatergic and cholinergic cells’ distribution was substantially overlapping for most of the parameters, as shown in the Raincloud analysis (Figure S4C).

Detailed analysis of the basal activity of optotagged cells, revealed remarkable heterogeneity within each neuronal population (Figure S5B-C). In further agreement with the heterogenous functional landscape of the LDT, using only optotagged cells, we identified 3 cholinergic, 4 glutamatergic and 3 GABAergic subclusters within each neuronal type (Figure S5D).

Next, we evaluated how pre-exposure to cocaine would trigger adaptations in these clusters. For this, ChAT-cre VGlut2-cre or vGAT-cre animals were injected with a cre-dependent AVV5-DIO-ChR2-eYFP in the LDT to allow optotagging. Animals were exposed to daily injection of cocaine for 5 days (COC); or vehicle (VEH). On the 6^th^ day, we recorded baseline activity and after a subsequent cocaine challenge, as previously described (Figure 2), followed by optotagging at the end of the electrophysiological recording, to preclude alterations in the activity of LDT neurons due to this stimulation protocol.

Pre-exposure to cocaine triggers significant differences in several electrophysiological characteristics of LDT neurons (Figure S6). Remarkably, this exposure alters the composition of LDT cholinergic, glutamatergic and GABAergic clusters (Figure S5E), which limits the direct comparison of VEH and COC neuronal clusters, as we are unable to track their initial cluster assignment. For this reason, we decided to restrict our analyses to optotagged LDT cells (see section below).

In summary, our data shows that LDT cholinergic, GABAergic and glutamatergic populations are functionally diverse, and that the segregation into three main subpopulations needs to be revised to include the different subclusters that we uncovered. This is essential in light of the distinct impact of cocaine pre-exposure on LDT neuronal subpopulations.

### Adaptive changes in cholinergic, glutamatergic and GABAergic LDT cells in cocaine pre-exposed animals

Our findings revealed that cocaine pre-exposure altered LDT electrophysiological signature and remodels neuronal clusters composition. For this reason, to better understand the differential impact of cocaine in each neuronal subtype, we decided to restrict our analyses to optotagged LDT cells, rather than per cluster analysis (Figure 3A; example of optotagged cells in Figure 3B).

**Figure 3.**
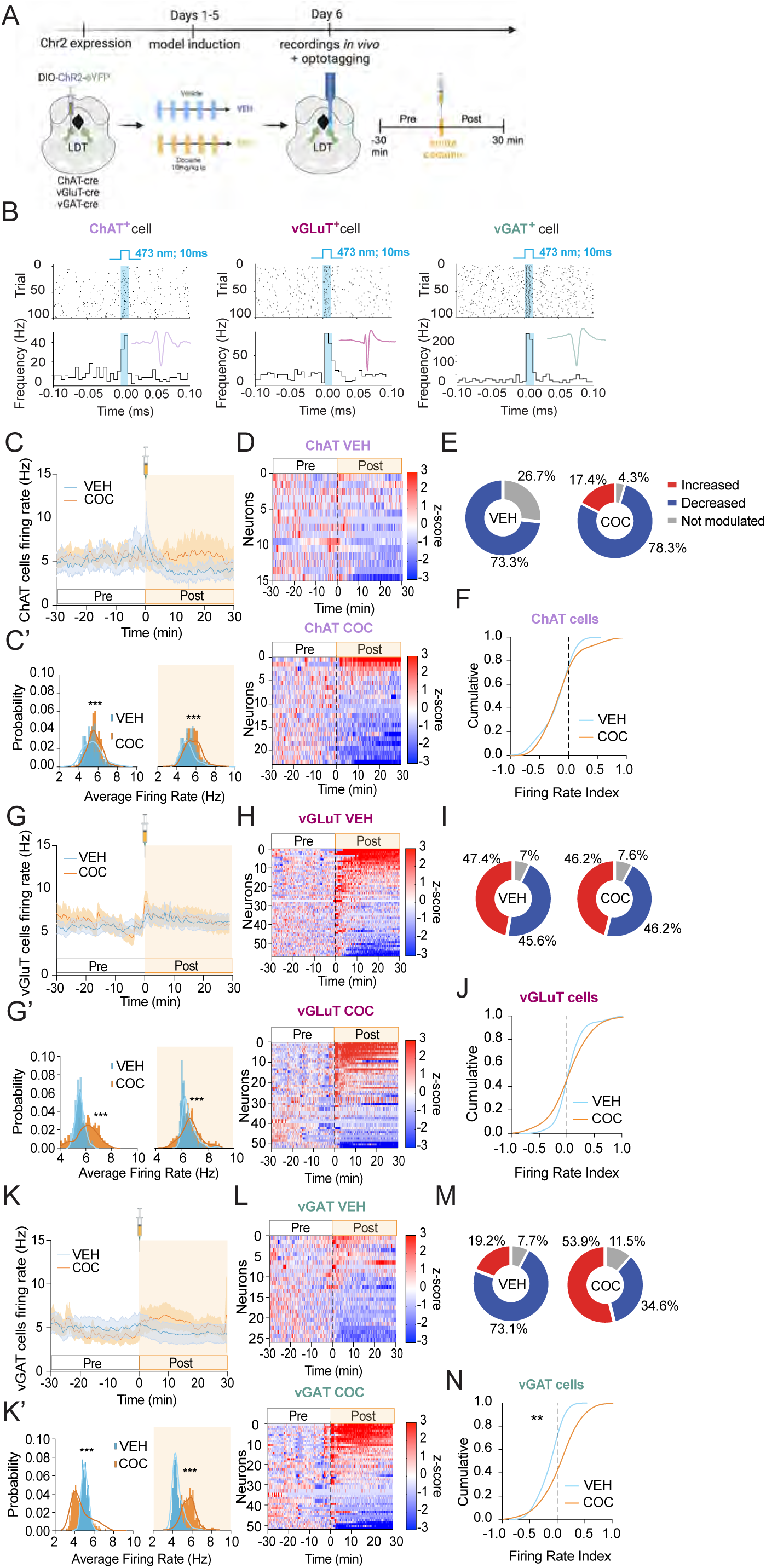
Cocaine pre-exposure differentially impacts distinct LDT neuronal populations. **A)** Schematic representation of the experimental design. ChAT, vGluT2 or vGAT-cre animals were injected with a cre-dependent ChR2 in the LDT. After opsin expression (4-6 weeks), animals were exposed to one daily injection of cocaine for 5 days (COC) or vehicle (VEH). On the 6^th^ day, animals were recorded in the LDT before and after acute cocaine injection. Optotagging protocol was performed in the end of the recordings. **B)** Examples of different optotagged cells. Top: Raster plot showing optotagged cell. Bottom: Frequency (Hz) during stimulation and baseline periods. **C)** PSTH depicting average firing rate of cholinergic neurons for COC (orange) and VEH (blue) groups (n_VEH-ChAT_ = 15, n_COC-ChAT_ = 23). We observe a shift to the right in the distribution of average firing rate in COC group in basal conditions (**C’**). Cocaine decreases LDT net activity in VEH group but this effect is attenuated in COC group. **D)** Heatmaps showing individual responses of ChAT^+^ cells to acute cocaine injection. **E)** Pie charts with % of neurons that are inhibited, excited or that present no change in activity in response to acute cocaine injection. Most of the cholinergic neurons of both groups are inhibited in response to acute cocaine, but COC group presents 17% of neurons that are excited. **F)** Distribution of the firing rate index in VEH and COC groups. No significant differences were found. **G)** PSTH depicting average firing rate of glutamatergic neurons (n_VEH-glut_ = 57, n_COC-glut_ = 52). We observe a shift to the right in the distribution of average firing rate in COC group in basal conditions (**G’**). (**H)** Heatmaps showing individual responses of vGluT2^+^ cells to acute cocaine injection. **I)** Pie charts showing that the response of glutamatergic neurons to acute cocaine injection is similar between groups. **J)** Distribution of the firing rate index of glutamatergic neurons in VEH and COC groups. No significant differences were found. **K)** PSTH depicting average firing rate of GABAergic neurons (n_VEH-GABA_ = 26, n_COC-GABA_ = 26). Contrary to cholinergic and glutamatergic neurons, we observe a shift to the left in the distribution of average firing rate in COC group in basal conditions (**K’**). Conversely, after acute cocaine, the distribution of COC group is shifted to the right, suggesting a more robust impact of acute cocaine in COC group. **L)** Heatmaps showing individual responses of vGAT^+^ cells to acute cocaine injection. **M)** The majority of GABAergic neurons of are inhibited in VEH animals, contrary to COC group, where 54% are excited. This is further confirmed by the shift in the **N)** firing rate index curve in comparison to VEH group. PSTH data represented as mean±SEM. *p < 0.05, **p < 0.01, ***p < 0.001.

We plotted the average firing rate of optotagged cells for each neuronal type in order to observe the net effect of cocaine pre-exposure. In baseline conditions, the distribution of the average firing rate of cholinergic cells was significantly different between COC and VEH groups, with COC group presenting increased average firing rate (Figure 3C-C’; Pre: Mann-Whitney, p=0.0096; average baseline firing rate of COC = 5.61 Hz, VEH = 5.34 Hz). Acute cocaine challenge decreases the average firing rate activity of cholinergic cells in VEH group, and this effect was attenuated in COC group (Figure 3C-D; VEH: Mann-Whitney, p < 0.0001; COC: Mann-Whitney, p = 0.7675). In COC group, 17% of cells increase activity in response to acute cocaine, which was not observed in VEH group (Figure 3D-E). Despite this, the firing rate index was not statistically significant between groups (Figure 3F; Mann-Whitney, p = 0.9286).

Regarding glutamatergic cells, in baseline conditions, the average firing rate was significantly higher in COC group (Figure 3G-G’; Pre: Mann-Whitney, p < 0,0001; average baseline firing rate of COC = 6.05 Hz, VEH = 5.54 Hz). Cocaine challenge increases average firing rate in both groups (Figure 4G-H, VEH and COC: Mann-Whitney, p < 0.0001); though half of the cells present excitations and other half inhibitions (Figure 4I). No differences between groups were found in the firing rate index of glutamatergic cells (Figure 3J; Mann-Whitney, p = 0.6339).

**Figure 4.**
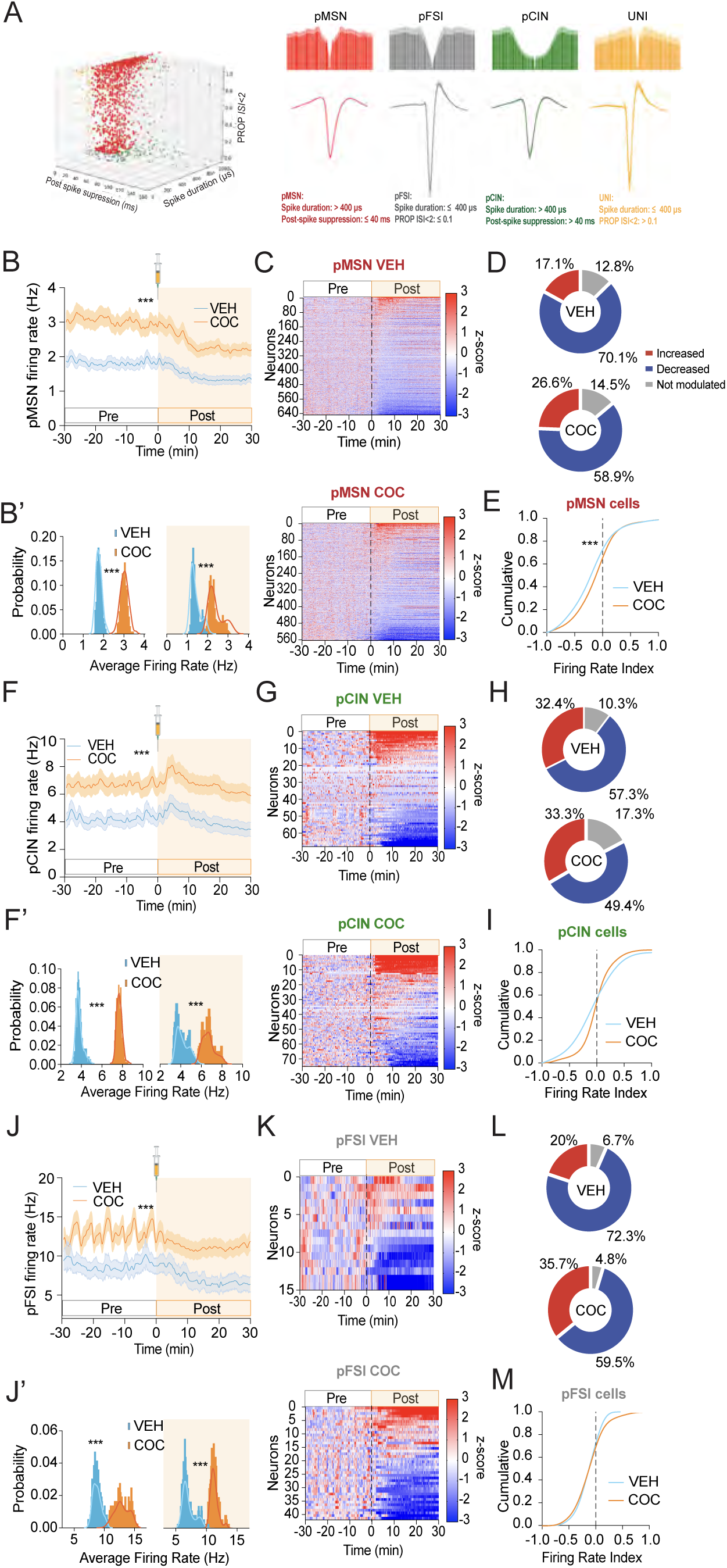
Cocaine-triggers neuronal adaptions in NAc neurons. **A)** Examples of waveforms and auto-correlogram of putative Medium Spiny Neurons (pMSN), Cholinergic Interneurons (pCIN), Fast Spiking interneurons (pFSI), and unidentified interneurons (unid). Classification of these neuronal populations is based on waveform duration (µS), post-spike suppression (ms), and proportion of the interspike intervals (PROPISI > 5s). **B)** PSTH depicting average firing rate of pMSNs for COC (orange) and VEH (blue) groups (n_VEH-MSN_ = 543, n_COC-MSN_ = 648). We observe a shift to the right in the distribution of average firing rate in COC group in basal conditions (**B’**). Acute cocaine decreases net activity of the NAc in both groups. **C)** Heatmaps showing individual responses of pMSNs to acute cocaine injection. **D)** Pie charts with % of NAc pMSNs that are inhibited, excited or that present no change in activity in response to acute cocaine injection. Most of the pMSNs are inhibited in response to acute cocaine, but COC group presents more neurons that are excited in relation to VEH. **E)** Distribution of the firing rate index of pMSN, showing a shift to the right in COC group. **F)** PSTH depicting average firing rate of pCIN (n_VEH-CINs_ = 75, n_COC-CINs_ = 68). We observe a shift to the right in the distribution of average firing rate in COC group in basal conditions (**F’**). After acute cocaine injection, there is an increase in activity during the first 10 min and then activity decreases to baseline levels in both groups. **G)** Heatmaps showing individual responses of pCIN to acute cocaine injection. **H)** Pie charts showing responses to acute cocaine injection is similar between groups. **I)** Distribution of the firing rate index of cholinergic neurons is not different between groups. **J)** PSTH depicting average firing rate of pFSI (n_VEH-FS_ = 15, n_COC-FSI_= 42). The average activity of pFSI is higher in COC group (**J’**). After acute cocaine, the activity of pFSI decreases in both groups, as confirmed in the **K)** heatmaps and **L)** pie charts. **M)** No differences in pFSI firing rate index curve were found between groups. PSTH data represented as mean±SEM. *p < 0.05, **p < 0.01, ***p < 0.001.

The increase in the firing rate of cholinergic and glutamatergic cells in COC group in baseline conditions can explain the net increase in LDT firing rate observed in Fig. 2C, as together, they comprise the majority of neurons in this region^9,32^.

Conversely, the average firing rate of GABAergic cells was significantly lower in COC groups (Figure 3K-K’; Mann-Whitney U= 55396.0, p < 0.0001; average baseline firing rate of COC = 4.50 Hz, VEH = 5.62 Hz). Interestingly, the effects of an acute cocaine injection were divergent in GABA cells from VEH and COC animals (VEH: Mann-Whitney, p < 0.0001; COC: Mann-Whitney, p < 0.0001). COC group cells present a robust increase in activity after acute cocaine injection, with 54% of cells increasing activity *vs* 19% of VEH group. VEH group present mostly inhibitions (73% of cells) in response to cocaine (Figure 3L-M). In agreement, the firing rate index of GABAergic cells was shifted to the right in COC group (Figure 3N; Mann-Whitney, p = 0.0113).

Overall, these results show that pre-exposure to cocaine triggers profound remodeling of the three neuronal populations of the LDT, that present changes in baseline activity and differentially respond to an acute cocaine challenge.

### Impact of cocaine pre-exposure in the activity of distinct NAc neuronal populations

Next, we aimed to understand the impact of pre-exposure to cocaine in different NAc neuronal populations. For this, we used established criteria to identify putative medium spiny neurons (pMSNs), cholinergic interneurons (pCINs) and fast spiking (pFS) interneurons (Figure 4A). Of 1468 recorded units, we identified 1211 pMSNs (n_MSNs-VEH_=648; n_MSNs-COC_=563), 143 pCINs (n_CINs-VEH_=68; n_CINs-COC_=75), 57 pFSs (n_FS-VEH_=15; n_FS-COC_=42), and 57 unidentified units (Figure 4A). The distribution of these neuronal types matched what has been described for this brain region^33^.

Interestingly, COC animals presented increased firing rate of pMSNs in comparison to VEH in baseline conditions (Figure 4B-B’; Pre: Mann-Whitney, p < 0.0001; average baseline firing rate of COC = 3.03 Hz, VEH = 1.82 Hz). Acute cocaine challenge decreased the activity of pMSNs in both groups, though this effect appeared more pronounced in COC group (Figure 4B; VEH: Mann-Whitney, p < 0.0001; COC: Mann-Whitney, p < 0.0001). The analysis of individual units showed that the majority of pMSNs were inhibited in response to cocaine (Figure 4C-D; 70% in VEH *vs* 59% in COC group), though there were also pMSNs that were excited in both groups (Figure 4D). The firing rate index of pMSNs was shifted to the right in COC group (Figure 5E, Mann-Whitney, p < 0.0001).

**Figure 5.**
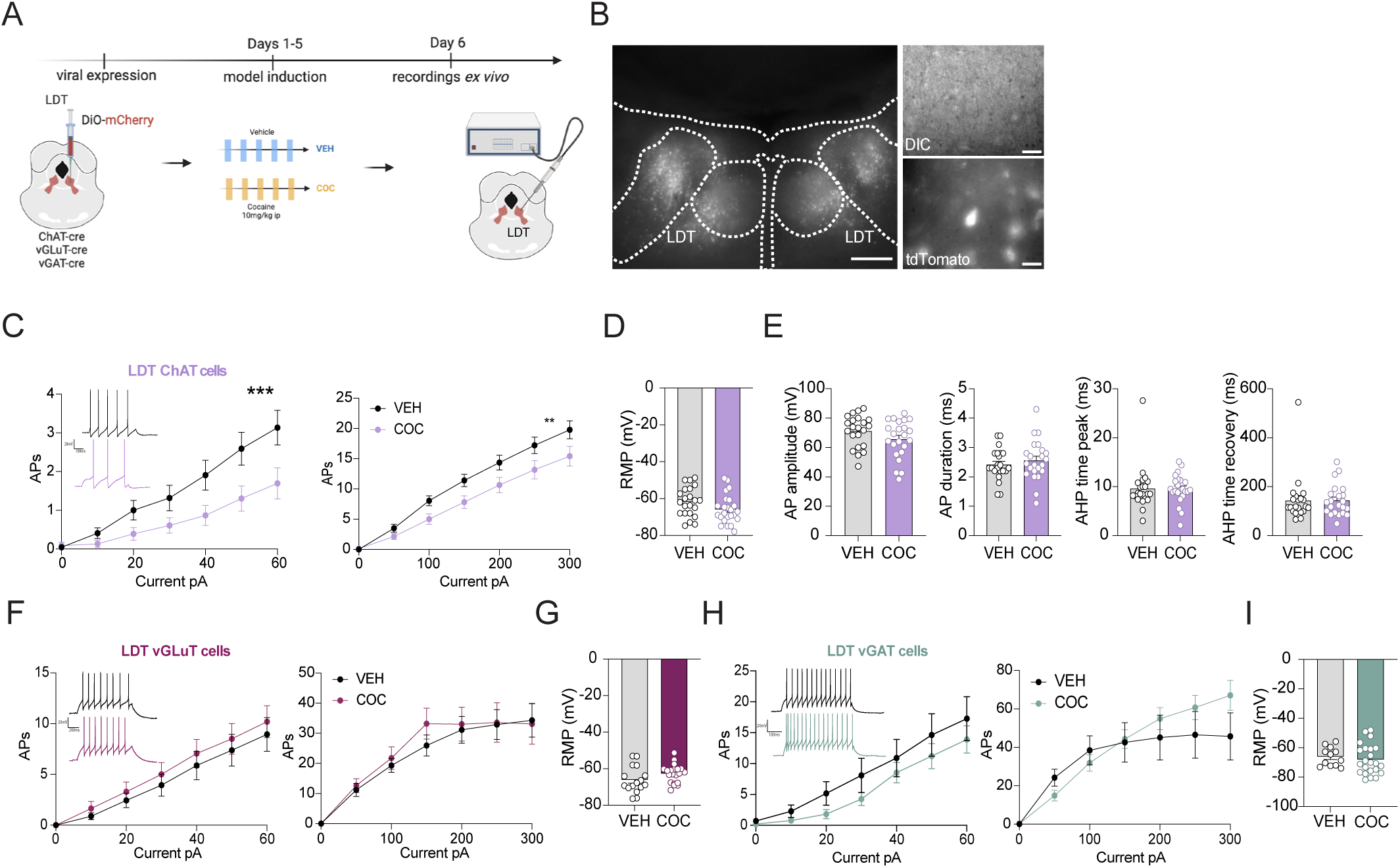
Pre-exposure to cocaine selectively alters LDT cholinergic cells. **A**) Schematic experimental time line to perform patch-clamp recordings in virally-tagged LDT neuronal populations after repeated vehicle (VEH) or cocaine (COC) injections. Dio-mCherry was injected in wither ChAT-cre, vGlut2-cre or vGAT-cre mice to allow tagging of specific LDT cell types. **B**) Image showing representative injection site of LDT glutamatergic neurons fluorescently tagged with mCherry. Visualization of representative mCherry-tagged GABA neurons in the electrophysiological set-up during patch-clamp recordings. **C**) Cocaine pre-exposure increased excitability to increasing current steps in LDT cholinergic neurons (number of cell/mice: saline n = 22/5 and cocaine n = 24/6). Increasing high current steps in LDT cholinergic neurons decreased excitability of cocaine pre-exposed neurons (\*\**P*=0.0020, 2way ANOVA) (number of cell/mice: saline n = 22/5 and cocaine n = 24/6). **D**) No significant changes were observed in RMP, **E)** AP amplitude, AP duration, AHP time peak and AHP time recovery (Unpaired t-test). **F)** No significant differences were observed in current curves of glutamatergic (number of cell/mice: saline n = 16/4 and cocaine n = 20/4) or **H)** GABAergic neurons (number of cell/mice: saline n = 12/3 and cocaine n = 23/4) nor in **G,I)** RMP. Representative voltage traces shown in insets for each condition. Data represented as mean±SEM. *p < 0.05, **p < 0.01, ***p < 0.001.

Regarding pCINs, COC animals presented increased firing rate in comparison to VEH in baseline conditions (Figure 4F-F’; Pre: Mann-Whitney, p < 0.0001). Despite this basal difference, the response to acute cocaine in VEH animals was significantly higher than what we observe in COC animals (Figure 4F-H; pre vs post: VEH: Mann-Whitney U = 80931.0, p < 0.0001; COC: Mann-Whitney, p = 0.6050). No significant differences between groups were found in the firing rate index, though there is a trend (Figure 4I; Mann-Whitney, p = 0.0884).

Regarding pFS neurons, COC animals presented increased firing rate in comparison to VEH in baseline conditions (Figure 4J-J’; Pre: Mann-Whitney, p < 0.0001). Cocaine injection decreased the activity of pFS in both groups (Figure 4J-L; VEH: Mann-Whitney, p < 0.0001; COC: Mann-Whitney, p < 0.0001), with no major differences between their firing rate index (Figure 4M; Mann-Whitney, p = 0.3813).

Overall, cocaine pre-exposure triggers persistent changes in the basal activity of putative MSNs, CINs and FSs, and alters their subsequent response to an acute cocaine challenge.

### Pre-exposure to cocaine selectively potentiates LDT-NAc cholinergic inputs

Our *in vivo* electrophysiological data showed that the LDT was robustly altered by repeated cocaine exposure, and that this triggered a differential response to a cocaine challenge. Moreover, cholinergic, GABAergic and glutamatergic neurons are differentially affected by cocaine pre-exposure. To further gain insights into the cellular adaptations induced by cocaine treatment, we performed patch-clamp recordings of cholinergic, GABAergic and glutamatergic neurons of the LDT in VEH and COC animals. To do this, we injected AAV-hSyn-Flex-mCherry in the LDT of ChAT-cre, vGluT2-cre or vGAT-cre transgenic mouse lines to selectively label and record mCherry^+^ cells (Figure 5A-B).

Repeated cocaine exposure did not significantly alter the resting membrane potential (RMP) of any of the LDT cell types recorded (Figure 5D, 5G, 5I). However, in COC mice, LDT cholinergic neurons responded with lower discharge frequency to depolarizing current injections in comparison to VEH group (Figure 5C; 2way ANOVA, p = 0.0001 and 2way ANOVA, p = 0.002). We did not find any significant differences in other parameters such as after hyper-polarization (AHP), recovery time, or AHP time peak, and action potential (AP) duration or amplitude (Figure 5E).

The change in firing activity was cell-type specific, as in striking contrast, glutamatergic and GABAergic LDT neurons exhibited comparable excitability curve regardless of the pre-treatment (Figure 5F; vGLuT: 2way ANOVA, p = 0.9776; Figure 5H, vGAT: 2way ANOVA, p = 0.8287).

The observed phenotypic heterogeneity of each LDT neuronal population (Figures S3-S5), could represent specific neuronal clusters that either receive and/or project to distinct downstream target regions, as it has been observed for VTA dopamine neurons, for example^34^. Hence, we next performed patch clamp recordings of the different LDT cell types in a projection-specific manner. This was achieved by injecting a retroAAV-FLEX-tdTomato in the NAc of ChAT-cre, vGluT2-cre and vGAT-cre animals to selectively record cholinergic, glutamatergic and GABAergic LDT-NAc neurons, respectively (ChAT-cre in Figure 6A-F; vGluT2-cre and vGAT-cre in Figure S7A-D). Remarkably, and in contrast to what we observed for all LDT cholinergic neurons (regardless of their projections), pre-exposure to cocaine triggered a marked upward shift of the excitability curve of cholinergic LDT-NAc projecting neurons (Figure 6C, 2way ANOVA, p = 0.0304) and a significant decrease in RMP (Figure 6D, unpaired t-test, p = 0.0354). These findings propose that LDT-NAc projecting cholinergic cells are more excitable, which is in line with the *in vivo* electrophysiological recordings (Fig. 3C).

**Figure 6.**
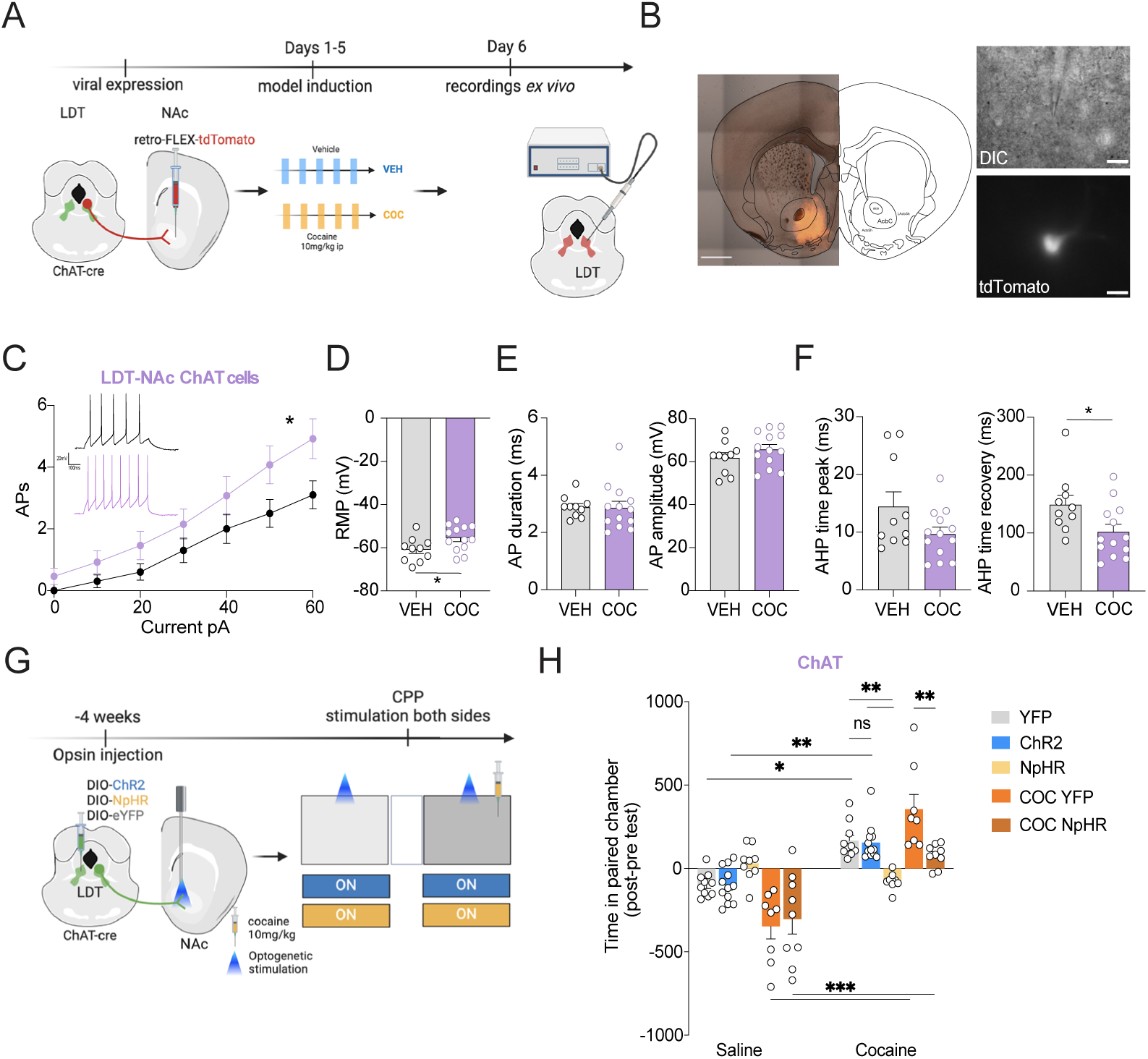
LDT-NAc cholinergic inputs are asymmetrically modulated by prior cocaine exposure and are required for cocaine conditioning. **A)** Schematic experimental time line used to perform patch-clamp recordings of LDT-NAc projecting neurons. A retro-FLEX-tdTomato was injected in the NAc, and red cells were recorded in the LDT. **B**) Example of retrograde virus injection in the NAc in ChATCre mice (left) and visualization of a cholinergic LDT-NAc neuron in the LDT (right). **C)** Cocaine pre-exposure significantly increased excitability and **D)** RMP of cholinergic LDT-NAc neurons (number of cell/mice: saline n = 10/5 and cocaine n = 13/8). **E**) Action potential duration and amplitude are not significantly affected by pre-exposure to cocaine. **F)** Pre-exposure to cocaine reduces the time to peak duration and recovery time of AHP in LDT-NAc cholinergic neurons. **G)** Schematic representation of the optogenetic manipulation experiments. ChAT-cre animals were injected in the LDT with a cre-dependent ChR2 for excitation, or NpHR for inhibition or YFP as control. COC animals were injected with YFP or NpHR. An optic fiber was placed in the NAc to allow stimulation of LDT terminals in this region. After opsin expression, optogenetic stimulation was performed in both sides of the apparatus and animals were conditioned to cocaine (10 mg/kg) to one side of the apparatus; saline was given to the other side. **H)** Optical activation of LDT-NAc cholinergic terminals had no effect on the observed preference for the cocaine-paired chamber, as ChAT-ChR2 animals were similar to YFP animals in terms of preference score (n_ChAT-ChR2_=11; n_ChAT-NpHR_=9, n_ChAT-YFP_=10). However, inhibition of LDT-NAc cholinergic neurons significantly reduced the preference for the cocaine-paired chamber. COC animals displayed a higher preference for the cocaine-paired chamber. This preference was reduced in animals with inhibition of LDT-NAc cholinergic terminals (n_COC-NpHR_=8, n_COC-YFP_=9). Data represented as mean±SEM. *p < 0.05, **p < 0.01, ***p < 0.001.

To further analyze changes occurring at cholinergic LDT-NAc projecting neurons, we examined several action potential characteristics. While the duration and amplitude remained unaltered (Figure 6E, duration: unpaired t-test, p = 0.9407; amplitude: unpaired t-test, p = 0.2422), we found that the AHP recovery time was significantly lower for the COC group than for the VEH group (Figure 6F; unpaired t-test, p = 0.0319). The same trend was seen for AHP time peak, although this difference is not significant (Figure 6F, unpaired t-test, p = 0.0767). Next, in ChAT-cre mice with selective labeling of cholinergic LDT-NAc neurons, we also recorded non-tagged LDT cholinergic neurons identified based on their shape and electrophysiological properties^35^. These putatively-non-NAc-projecting cholinergic cells exhibited a decrease in their excitability profile (confirming data shown in Figure 5C).

When examining the input-output responsiveness of either glutamatergic or GABAergic LDT-NAc neurons, we did not find any major effect of cocaine pre-exposure when compared to VEH mice (Figure S7C-D).

Collectively, these results reveal an impact of cocaine pre-exposure on LDT-NAc neurons, with selective and asymmetric adaptations occurring at distinct cholinergic cells of the LDT.

### Inhibition of LDT-NAc cholinergic inputs blunts cocaine-induced CPP

Our data showed that LDT-NAc cholinergic cells were altered due to repeated cocaine pre-exposure, indicating that this pathway could be an important target for the modulation of cocaine’s behavioral effects. Thus, we conducted a series of conditioned place preference (CPP) experiments combining cocaine administration with optogenetic manipulation of LDT terminals in the NAc. Expression of virus and optical fiber placement was checked for all the subjects (Figure S8). Animals were conditioned using a standard CPP protocol using cocaine (10 mg/kg) as reinforcer, and optogenetic stimulation was used to either activate or inhibit LDT terminals expressing ChR2 or NpHR, respectively (Figure 6G; Figure S9). We included 5 groups of animals: naïve animals injected with YFP (control for stimulation), or ChR2 or 3 NpHR, and COC animals either injected with YFP or NpHR. Considering that optical stimulation of LDT-NAc cholinergic inputs *per se* trigger preference^4^, we stimulated LDT-NAc terminals in both sides of the CPP apparatus to exclude any confounding effects of the stimulation alone. One of the chambers was paired with cocaine and the other side with saline; the order and side was counterbalanced between animals.

Optical activation of LDT-NAc cholinergic terminals had no effect on the observed preference for the cocaine-paired chamber, as ChAT-ChR2 animals were similar to YFP animals in terms of preference score (Figure 6H; 2way ANOVA F(4, 43) = 10.31, p<0.0001, Bonferroni post-hoc comparison, p=0.9998). Conversely, inhibition of LDT cholinergic neurons significantly reduced the preference for the cocaine-paired chamber in naïve animals (Figure 6H; Bonferroni post-hoc comparison, p=0.005).

Interestingly, COC mice presented a trend for higher preference for the cocaine-paired chamber (vs YFP, Bonferroni post-hoc comparison, p=0.061). In line with the data of naïve animals, inhibition of LDT-NAc cholinergic terminals significantly reduced time spent in the cocaine-paired chamber when compared to COC YFP mice (Bonferroni post-hoc comparison, p=0.0019).

As control experiments, we also modulated the activity of glutamatergic and GABAergic LDT-NAc terminals. Activation of glutamatergic LDT-NAc projections did not trigger significant changes in the time in the cocaine-paired chamber in the CPP between groups; though inhibition of these projections reduced place preference for the cocaine-paired chamber in the CPP (Figure S9C). Modulation of LDT-NAc GABAergic inputs did not impact cocaine place preference (Figure S9D).

Our previous data showed that LDT-NAc cholinergic neurons were essential in mediating cocaine reinforcing properties. To validate and extend these findings, we designed another experiment, in which cocaine was given in the two sides of the apparatus to make them equally preferred to the animal, while optical stimulation was only applied to one of the sides (Figure S9A). In further support of the necessity of LDT-NAc cholinergic inputs in mediating cocaine rewarding effects, inhibition of LDT cholinergic neurons during cocaine conditioning significantly decreased time spent in the stimulated chamber (Bonferroni post-hoc comparison, p<0.0001). In turn, optical activation of LDT-NAc cholinergic terminals resulted in significantly more time spent in the stimulated chamber compared to the non-stimulated side (Figure S9E; 2way ANOVA F (2, 27) = 32.19, p<0.0001; Bonferroni post-hoc comparison, p<0.0001). In this task, optical modulation of glutamatergic LDT-NAc inputs did not affect preference for either chamber (Figure S9F). Inhibition of GABAergic LDT-NAc inputs slightly decreased preference for the side with cocaine and stimulation (Figure S9G; 2way ANOVA F(2, 25) = 4.004, p = 0.0310; Bonferroni post-hoc comparison, p = 0.0005).

In summary, these results indicate modest effects of manipulating glutamatergic and GABAergic LDT-NAc inputs in cocaine conditioning. On the other hand, activation of LDT cholinergic neurons robustly enhances the motivational value of the cocaine-paired environment, and suppression of LDT-NAc cholinergic activity robustly dampens the reinforcing properties of cocaine.

## Discussion

By integrating *in vivo* dense silicon probe recordings, optotagging and patch-clamp electrophysiology, we provide unprecedented insights into the circuit-specific neuroadaptations induced by cocaine pre-exposure in the LDT-NAc circuit. Our findings reveal that cocaine pre-exposure triggers persistent neuroplastic adaptations in the LDT-NAc circuit, selectively modulating distinct neuronal populations, and triggering a projection-specific effect in LDT cholinergic cells. Furthermore, we provide evidence that LDT-NAc cholinergic signaling plays a crucial role in cocaine-induced conditioning.

### Cocaine exposure induces LDT-NAc circuit-specific adaptations

Our data shows that LDT cholinergic, glutamatergic and GABAergic cells project to the NAc, and overall, LDT neurons present a preferentially innervation of MSNs and CINs, extending previous knowledge about LDT projections to the NAc ^4,10,36^.

A key finding is that repeated cocaine exposure robustly alters the baseline activity and acute cocaine-evoked responses in both the LDT and NAc. The increased baseline firing rate observed in both regions after cocaine pre-exposure suggests the appearance of persistent neuroadaptations. Though it was not the main objective of this work, we found enduring LDT alterations 15 days after the last injection of cocaine, which could significantly influence how this region responds to subsequent cocaine challenges even after withdrawal periods.

Our *in vivo* data aligns with previous *ex vivo* studies showing that repeated cocaine exposure triggers pre- and post-synaptic plasticity changes in the LDT ^20,21,37^, and also in NAc neurons ^22,23,38^. Alterations in the LDT could influence modulatory signals in downstream brain regions, and ultimately, cocaine-associated behavioral effects. For example, intra-VTA injection of muscarinic or nicotinic acetylcholine receptors antagonists before cocaine conditioning suppresses cocaine CPP development, suggesting that the activity of LDT cholinergic neurons (main cholinergic source in the VTA) contribute to the acquisition of the rewarding effects of cocaine ^39^. In addition to the canonical VTA, we now add a new player to the cocaine circuitry, as our RRR analysis reveals that cocaine pre-exposure significantly modifies the direct functional connectivity between LDT and NAc.

### Functional heterogeneity within LDT neuronal populations

A striking finding was the observed functional diversity within cholinergic, GABAergic, and glutamatergic LDT neurons. Despite sophisticated clustering approaches, we were unable to definitely categorize neurons into genetically-defined clusters. This stands in contrast to other brain regions where distinct neuronal populations often exhibit characteristic firing patterns and waveform properties, such as the striatum (this work;^40^) or hippocampus^41^. The overlap in multiple electrophysiological parameters of different optotagged cell types, and the existence of multiple subclusters identified within each cell type, demonstrates a previously unappreciated functional diversity within the LDT. Notably, cocaine pre-exposure altered the electrophysiological properties of LDT neurons and modified cluster distribution, highlighting the dynamic nature of these cellular properties. These observations underscore the importance of combining multiple technical approaches, including optical tagging and projection-specific targeting, to reliably identify and characterize distinct neuronal populations in this region in subsequent studies.

### Cell-type specific effects of cocaine pre-exposure in the LDT

Using genetically-identified cells as ground-truth data, we were able to attest that cocaine pre-exposure triggers robust plasticity changes in the three main LDT neuronal subtypes *in vivo and ex vivo*.

*In vivo*, COC group display slightly increased firing rate of LDT cholinergic neurons in baseline conditions, and present a higher number of neurons that were excited in response to an acute cocaine injection, in relation to VEH group. This is in line with *ex vivo* studies in rats reporting that cocaine pre-exposure increased excitatory synaptic transmission to LDT cholinergic neurons due to a presynaptic plasticity mechanism^37^ and/or the induction of intrinsic membrane plasticity, resulting in increased firing activity of a subset of LDT neurons^20^.

Importantly, our *ex vivo* recordings revealed a surprising asymmetrical effect of cocaine pre-exposure in LDT-NAc-projecting cholinergic neurons *vs* non-NAc-projecting cells. Though we cannot fully exclude that some unlabeled cells can be non-infected LDT-NAc-projecting neurons, the probability is low, considering the sparse number of these inputs (Figure 1). This unequal effect of repeated cocaine exposure in LDT-NAc-projecting-cholinergic cells vs all-LDT-cholinergic cells can pinpoint to parallel actions of these different neurons in reward and aversion circuitry. For example, we have shown that activation of LDT-NAc cholinergic inputs change natural rewards value and shifts preference (rewarding), whereas LDT-VTA activation induced strong freezing responses^5,42^.

The selective potentiation of LDT-NAc cholinergic pathway may represent a key mechanism through which cocaine pre-exposure can impinge changes in the reward circuit to contribute for addictive behaviors. In line, our *in vivo* recordings revealed that while acute cocaine typically suppressed net cholinergic neuron firing in drug-*naive* animals, this response was notably different in animals previously receiving repeated cocaine injections. This adaptation may reflect the development of tolerance at the cellular level, similar to what has been observed in other reward-related circuits^38^. The enhanced excitability in NAc-projecting cholinergic neurons also aligns with studies showing that cocaine exposure increases acetylcholine release in the NAc^43–45^, though it remains to be determined if this acetylcholine arises from the LDT and/or local CINs.

### LDT-NAc cholinergic inputs modulate cocaine conditioning

The observed enhanced cholinergic signaling may be particularly important for the processing of drug-associated cues/contexts, as previous work has shown that NAc cholinergic signaling is crucial for associative learning^46–49^. Interestingly, we found that LDT-NAc cholinergic inputs activation enhances cocaine CPP, whereas inhibition attenuates it in both naïve and COC animals, which demonstrates that this pathway plays a crucial role in the assignment of value to a cocaine-associated context. This extends previous work showing that LDT cholinergic signaling contributes to natural reward processing^4,15^ and suggests that prior exposure to repeated cocaine may hijack this circuit to promote drug-seeking behavior.

Initial studies using pharmacological inhibition of cholinergic LDT neurons showed attenuation of cocaine-induced CPP, leading to the interpretation that afferents to the VTA were the main actors of this phenomenon^39^. Now, we add a new player to the equation, and the observed divergent effects on NAc-projecting versus non-projecting cholinergic neurons may even help reconcile seemingly contradictory findings in the literature. For instance, a study found that selective ablation of LDT cholinergic neurons did not affect cocaine reward, but they reported reduced responsiveness to cocaine-associated cues^7^. Thus, global manipulation of LDT cholinergic neurons can fail to capture the specific role of LDT projection-specific subpopulations.

## Conclusions

Our findings establish the LDT-NAc cholinergic pathway as a key mediator of cocaine’s conditioning effects and reveal complex circuit cell-specific adaptations induced by repeated cocaine exposure. The asymmetric effect of repeated cocaine exposure in LDT-NAc projecting *vs* other LDT cells strongly suggests that in the LDT, there are subclasses of cholinergic neurons with distinct physiological properties and vulnerabilities to cocaine effects, as our *in vivo* data confirmed. These differences may arise from their anatomical specificities, i.e., where these cells project to and/or receive inputs from. Though we highlighted the contribution of LDT cholinergic cells for cocaine conditioning, we also found modest neuroplastic changes in GABAergic and glutamatergic subtypes. This reinforces the need to perform additional studies in order to understand how cocaine alters this brain region shaping different neuronal subtypes in a selective manner, and their contribution for cocaine-related behaviors.

## Methods

### Experimental animals and housing conditions

Wild type male and female mice (2-5 months old, C57BL6/J) were used. The progeny produced by homozygous vGAT-cre (Vgat-IRES-Cre: Slc32a1tm2(cre)Lowl/J, The Jackson Laboratory), vGluT-cre (VGluT2-IRES-Cre:Slc17a6tm2(cre)Lowl/J, The Jackson Laboratory), SOM-cre (Sst-IRES-Cre mice, JAX #013044) and PV-Cre (#8069, The Jackson Laboratory) transgenic mice were used. The progeny produced by mating ChAT-cre (ChAT-IRESCre: B6;129S6-Chattm2(cre)Lowl/J, #006410, The Jackson Laboratory), D1-cre (line EY262, Gensat.org) and D2-cre (line ER44, Gensat.org) heterozygous transgenic male mice with wild-type C57/Bl6 females were genotyped at weaning by PCR fragment analysis. All animals were maintained under standard laboratory conditions: artificial 12 h light/dark cycle, temperature of 21 ± 1°C and 60% relative humidity. Mice were given standard diet (4RF21, Mucedola, Italy) and water *ad libitum*. All behavioral experiments were performed during the light period of the light/dark cycle. Health monitoring was performed according to FELASA guidelines and procedures were conducted in accordance with European Union regulations (Directive 2010/63/EU). Animal facilities and the people directly involved in animal experiments received certification from the Portuguese regulatory entity - Direção Geral de Alimentação e Veterinária (DGAV). All protocols were approved by the Ethics Committee of the Life and Health Sciences Research Institute (ICVS) and by DGAV (approval reference #8332, dated 2021-05-08).

### Stereotaxic Surgery and Viral Vector Injection

Mice were anesthetized using sevoflurane (2%–3%, plus oxygen at 1-1.5 l/min) and placed in a stereotaxic frame (Kopf Instruments). Body temperature was maintained at 36°C using an animal temperature controller (ATC2000, World Precision Instruments). AAV viral vectors expressing channelrhodopsin (ChR2) under a cre-dependent promoter (AAV5-EF1a-DIO-hChR2(H134R)-eYFP) were injected unilaterally into the LDT using stereotaxic coordinates (AP: -5.2 mm, ML: 0.6 mm, DV: - 3.4 mm) for electrophysiological recordings. The injection volume was 300 nL per site delivered over 5 minutes using a Nanojet III Injector (Drummond Scientific, USA) at a rate of 1 nl per second. After, the needle was left in place for an additional 5 minutes to allow diffusion before slowly being retracted. Mice were allowed to recover for at least 4 weeks before exposure to cocaine or vehicle for electrophysiological recordings.

For *ex vivo* electrophysiological recordings, ChAT-Cre, vGAT-Cre and vGluT2-Cre mice respectively, were injected bilaterally in the LDTg (AP: -4.7 mm, ML: ± 0.5 mm, DV: − 3.6 mm) with 300nL of AAV8-DIO-mCherry (Addgene).To specifically identify neurons projecting to the NAc, an pAAVrg-FLEX-tdTomato (Addgene) was injected bilaterally in the NAc of ChAT-Cre, vGluT2-Cre and vGAT-Cre mice respectively. Only male mice were used and were 8 weeks old at the time of surgery. Experiments started 3 weeks after injections to allow sufficient viral expression and recovery from the surgery.

Animals used in conditioned place preference experiments were injected with 300nl of cre-dependent AAV5/EF1a-DIO-ChR2(H134R)-eYFP, AAV5/EF1a-DIO-eNpHR-eYFP, or AAV5/EF1a-DIO-eYFP (UNC vector core) into the LDT (as above). After the injection, a 0.2-mm-diameter optic fiber (Thorlabs) was slowly lowered into the right mouse NAc (AP: 1.4 mm, ML:0.9 mm, DV: -4.5 mm). Once in place, the fiber was secured to the skull using dental cement (Superbond C&B kit). At the end of the surgical procedure, mice were removed from the stereotaxic frame, placed back in their home-cages on a pre-heated pad at 37°C until complete recovery and subcutaneously injected with buprenorphine (0.05 mg.kg−1), as well as once every 24 h during three successive days. Animals were let to recover for 4 weeks before behavioral procedures.

### *In vivo* electrophysiological recordings and data acquisition

Neural electrical activity was recorded from two 64-channel silicon shank Neuronexus probe using a multichannel acquisition processor (Open Ephys Acquisition Board). Wide-band (0.1–8000 Hz) neurophysiological signals from the probe were amplified 1,000 times via the Intan RHD2164 series Amplifier System and were continuously acquired at 30 kHz. Recordings were performed concomitantly to optogenetic blue light (450 nm) stimulation for photo-identification of ChAT, VGAT and VGluT neurons (100 trials: 10 ms; every second performed at the end of the recording session). The software-controlled laser was connected and synchronized to the recordings via TTL pulses. The multichannel processor recorded simultaneously the laser pulses and the neuronal activity.

### Firing rate analysis

Spike sorting was performed semi-automatically using the clustering software Kilosort2 (https://github.com/MouseLand/Kilosort/tree/v2.0 and https://github.com/SjulsonLab/Kilosort2Wrapper) the graphical spike-sorting application phy (https://github.com/cortex-lab/phy). The data were then imported into Python for analysis using custom-written code.

Peri-event histograms (PSTH) were constructed for the period spanning 30 minutes before to 30 minutes after the acute cocaine injection, using 5-second bins. To assess the relative firing rate of cells during the Pre and Post intervals, we computed a firing rate index (ϕ) as follows:

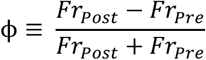

Where Fr_Pre_ is the firing rate of the cell during the Pre interval, and Fr_Post_ is the firing rate of the cell during the Post interval. This index provides a standardized measure of firing rate changes: A value of ϕ = 0 indicates equal firing rates during both intervals. A value of ϕ = -1 indicates that the cell fires exclusively during the Pre interval. A value of ϕ = 1 indicates that the cell fires exclusively during the Post interval.

A cell was classified as responsive through optotagging techniques if it tested positive in at least two out of three methodologies ^50–52^. This multi-method approach ensured a robust and reliable identification of opto-tagged cells, minimizing false positives.

Immediately after the recordings were concluded, the animals were injected with a lethal dose of mixture of ketamine/medetomidine and were transcardially perfused with 0.9% saline, followed by 4% paraformaldehyde (PFA) solution. The brains were extracted and processed for histology to confirm the recording locations. Histological confirmation of the probe location was achieved by applying DiI lipophilic carbocyanine dye (1%; Sigma-Aldrich) to the probes before insertion.

### *Ex vivo* electrophysiological recordings and data acquisition

On day 6, after 5 consecutive days of intraperitoneal injection of either saline (10ml/kg) or cocaine (10mg/kg), mice were anesthetized (Ketamine 150 mg/kg/Xylazine 10 mg/kg) and transcardially perfused with aCSF for slice preparation. Coronal 250 μm slices of LDTg were obtained in bubbled ice-cold 95% O2/5% CO2 aCSF containing (in mM): KCl 2.5, NaH2PO4 1.25, MgSO4 10, CaCl2 0.5, glucose 11, sucrose 234, NaHCO3 26.

Slices were then incubated in aCSF containing (in mM): NaCl 119, KCl 2.5, NaH2PO4 1.25, MgSO4 1.3, CaCl2 2.5, NaHCO3 26, glucose 11, at 37 °C for 15min, and then kept at room temperature.

Slices were transferred and kept at 32–34 °C in a recording chamber superfused with 2.5 ml/min aCSF. Visualized whole-cell current-clamp recording techniques were used to measure excitability, using an upright microscope (Olympus France).

Current-clamp experiments were obtained using a Multiclamp 700B (Molecular Devices, Sunnyvale, CA). Signals were collected and stored using a Digidata 1440 A converter and pCLAMP 10.2 software (Molecular Devices, CA). In all cases, access resistance was monitored by a step of − 10 mV (0.1 Hz) and experiments were discarded if the access resistance increased more than 20%. Internal solution contained (in mM): K-D-gluconate 135, NaCl 5, MgCl2 2, HEPES 10, EGTA 0.5, MgATP 2, NaGTP 0.4. Depolarizing (0–300 pA) or hyperpolarizing (0–450 pA) 800 ms current steps were used to assess excitability and membrane properties of LDTg neurons.

Offline analysis was performed using Clampfit 10.2 (Axon Instruments, U.S.A.) and Prism (Graphpad, U.S.A.).

### Behavioral tests

#### Open field – locomotor sensitization

Locomotor activity was evaluated using transparent open-field chambers (Med Associates), each equipped with a 32 × 32 photobeam array. The Med Associates system automatically recorded photobeam breaks as a measure of movement. Prior to testing, mice were acclimated to the experimental room for at least one hour. Following this habituation period, individual mice received an intraperitoneal injection of cocaine hydrochloride (10 mg/kg) and were reintroduced into the chamber. Cocaine-induced locomotor activity was recorded for 60 minutes. The collected locomotor parameters included total distance traveled (cm). Data were recorded in 5-minute intervals.

The cocaine dose and sensitization protocol selected (10 mg/kg) falls within the effective range (10–20 mg/kg) for inducing moderate to high levels of locomotor sensitization (Blanchard et al., 1998). A dose of 10 mg/kg was specifically selected in this study to minimize the risk of ceiling effects on behavioral responses (Partridge and Schenk, 1999).

Cocaine administration was repeated at the same time each day for five consecutive days. This was followed by a sixth-day electrophysiological recording session.

To assess the influence of repeated handling and injection procedures on locomotor behavior in the absence of cocaine, a group of animals (n = 10/group) underwent saline injection (VEH group).

#### Conditioned Place Preference (CPP)

One week before the optogenetic experiments, the animals were daily handled for 30 min, at least once a day, and habituated to being connected to the optic fiber. All tests are performed during the light phase of the light/dark cycle in dimly lit room. The behavioral setup was always cleaned with ethanol 10%, and distilled water, before each animal session. The CPP protocol was adapted from a previously published report. The CPP apparatus consisted of two compartments with different patterns, separated by a neutral area (Med Associates): a left chamber measuring 27.5 cm × 21 cm with black walls and a grid metal floor; a center chamber measuring 15.5 cm × 21 cm with gray walls and gray plastic floor; and a right chamber measuring 27.5 cm × 21 cm with white walls and a mesh metal floor.

Mice location within the apparatus during each preference test was monitored using a computerized photo-beam system (Med Associates). <u>Cocaine conditioning:</u> Briefly, on day 1, individual animals were placed in the center chamber and allowed to freely explore the entire apparatus for 20 min (pre-test). On days 2 and 3, mice were injected with saline 0.9% and confined to one side chamber for 30 min with optical stimulation (saline side); in the second session, mice were injected with cocaine (10 mg/kg, i.p.) and confined to one of the side chambers for 30 min and paired with optical stimulation, cocaine side. On day 4, mice were allowed to freely explore the entire apparatus for 20 min (post-test). Optical stimulation consisted of 80 pulses of 10 ms at 20 Hz, every 15 s with a blue laser and 4 s of constant light at 10mW with a yellow laser. Results are expressed as the difference of time spent on each side. <u>Stimulation conditioning:</u> On days 2 and 3, animals were injected with cocaine (10 mg/kg, i.p.) and confined to one of the side chambers for 30 min and paired with optical stimulation - Cocaine+stimulation; in the second session, mice were injected with cocaine (10 mg/kg, i.p.) and confined to the other side chamber for 30 min with no optical stimulation - Cocaine. Sessions were counterbalanced between animals.

#### Immunohistochemistry

Mice were deeply anesthetized with a mixture of ketamine/medetomidine and subjected to transcardial perfusion with saline 0.9%, followed by 4% paraformaldehyde (PFA) in PBS. Post-perfusion, brains were harvested and post-fixed in 4% PFA at 4°C overnight. Subsequently, brains were cryoprotected in 30% sucrose for 48–72 hours. Coronal sections (40 μm thick) were prepared using a vibratome (Leica) and stored in PBS with 0.01% sodium azide. Free-floating sections were washed in PBS three times (5 min each), then blocked for 1 hour in PBS containing 5% normal donkey serum and 0.3% Triton X-100. Sections were incubated overnight at room temperature with one or more primary antibodies diluted in blocking buffer: goat anti-ChAT (1:750; AB144P, Millipore), rabbit anti-GFP (1:1000; Abcam) or goat anti-GFP (1:1000, Abcam) for Channelrhodopsin-2(ChR2) amplification, rabbit anti-cFos (1:500; Millipore), chicken anti-mCherry (1:1000; HenBiotech). The following day, sections were rinsed in PBS three times (15 min each) and incubated with species-appropriate secondary antibodies (5 μg/ml) conjugated to AlexaFluor 488, 594, or 647 (Invitrogen) for 2 hours at room temperature. Afterward, sections were washed in PBS and mounted onto glass slides using Fluoromount (Invitrogen) with DAPI (0.5 μg/ml). Imaging was performed using a confocal microscope (Olympus FV3000).

#### RNA scope *in situ* hybridization

C57BL/6J wild-type mice were anesthetized with a mixture of ketamine/medetomidine and decapitated 60 mins after injection of cocaine (i.p. 10 mg/kg). The brains were rapidly extracted, flash-frozen in isopentane chilled on dry ice, and stored at −80°C. Cryostat sections (16 μm) were mounted onto glass slides and stored at −80°C until processed. For the assay, sections were fixed in 4% PFA for 15 minutes at 4°C, followed by dehydration through graded ethanol and treatment with protease IV. RNA Scope Multiplex Fluorescent Reagent Kit V2 (Advanced Cell Diagnostics, Hayward, CA; Cat. #323100) was used to perform RNA scope in situ hybridization for *c-fos, vGAT, vGLuT* and *ChAT* mRNA. Antisense RNA hybridization probes were used to detect transcripts encoding *c-fos* (c-fos; -C1), *vGLUT2* (Slc17a6; 319171-C2), *ChAT* (Chat; 408731-C4), and *VGAT* (Slc32a1; 319191-C3; Advanced Cell Diagnostics) during the hybridization step (2 h; 40 °C). After, sections were subjected fluorophore incubation with Opals 520, 570, 620 e 690 (Akoya Biosciences). After each step in the protocol, the sections were washed two times with 1X wash buffer provided in the kit. All slides were stained for 4’,6-Diamidino-2-phenylindole (DAPI), mounted with Prolong Gold Antifade mounting media and stored at 4 °C in the dark. The covered sections (Vectashield mounting medium; Vector Laboratories) were imaged and photographed with PhenoImager HT at final magnification of 20X.

### Clusterization

#### Cluster Analysis with WaveMAP

To identify neurons with waveform characteristics similar to our optotagged cells, we employed a waveform-based clustering approach. For this analysis, we extracted the mean waveform of each neuron and normalized all waveforms using their vector norm to eliminate amplitude-related bias. This normalization ensured that clustering was driven by waveform shape rather than absolute amplitude. The standardization protocol followed was based on WaveMAP. This tool was used to identify neuron clusters based on waveform characteristics through Uniform Manifold Approximation and Projection (UMAP) ^29^. After defining the clusters, we included optotagged cells in the UMAP representation to assess their distribution among the identified clusters.

#### Cluster Analysis with CellExplorer

As an alternative approach, we used CellExplorer, an open-source tool that computes standardized electrophysiological metrics and classifies neurons based on various functional properties, including spike waveform, spiking statistics, monosynaptic connections, and behavioral spiking dynamics, among other parameters^30^. We extracted 22 electrophysiological parameters, which were then exported to Python for analysis. To reduce data dimensionality while preserving most of the variance, we applied

Principal Component Analysis (PCA; sklearn.decomposition.PCA) to an N × D matrix, where N represents the number of cells and D corresponds to the extracted electrophysiological parameters. We retained principal components (PCs) that explained at least 80% of the variance in the dataset. The reduced data were then clustered using the K-means algorithm (sklearn.cluster.KMeans).After defining the clusters, we included optotagged cells to assess their distribution among the identified clusters.

#### Reduced rank regression model

To quantify how LDT population activity is related to NAc responses, we fit a reduced-rank regression (RRR) model via scikit-learn’s PLSRegression, treating LDT spike counts as the predictor matrix X and NAc spike counts as the response matrix Y. In this implementation the r components argument constrains the coefficient matrix B to rank r, imposing an implicit low-dimensional regularization: only r latent axes in X may influence Y. We determined the optimal r by five-fold cross-validation on each 30 min recording period (baseline and post-injection). For each candidate r between 1 and 20, we randomly partitioned 20 min of data into five equally sized folds, trained on four folds, and computed the held-out r-squared on the fifth. We repeated this across the five folds and selected the r that maximized mean out-of-fold r-squared. Consistently across animals and conditions, this procedure peaked at r = 3, which we then held fixed for all subsequent analyses.

For each animal and each condition (30 min pre-injection, 30 min post-injection), we binned spike times into non-overlapping 10 ms intervals to form two matrices: X (shape T x N_LDT) and Y (shape T x N_NAc), where T = 180,000 bins. From these T timepoints, we randomly selected 20 min for model fitting and reserved the remaining 10 min for held-out evaluation.

After fitting each RRR model, we extracted its coefficient matrix B and performed a singular-value decomposition (SVD), B = U Sigma V^T. We recorded the vector of singular values of Sigma and their sum as a scalar index approach of the model’s effective communication strength, i.e., the total covariance explained by the r latent modes.

To probe how stable the learned LDT to NAc mapping remains over time and in response to cocaine, we applied the baseline-trained RRR model (fitted to the first 5 min of pre-injection data) to sliding 5 min non-overlapping windows covering 30 min before and 30 min after injection. We then used the baseline RRR model to predict each neuron’s firing rate index from the corresponding LDT spike counts in one 5-minutes window and check the performance by r-squared metric.

### Statistical Analysis

All data are presented as mean ± SEM unless otherwise specified. Statistical analyses were performed with Prism 10.0 (GraphPad Software) or Python. All statistical tests were two-tailed, and results were considered statistically significant when the p value was less than 0.05. Normality and equal variances between group samples were assessed using the Kolmogorov-Smirnov normality test and Brown-Forsythe tests, respectively. When normality and equal variance between sample groups were achieved, the paired t test, unpaired t test, one-way ANOVA or two-way repeated-measures ANOVA with multiple comparisons was used. When normality or equal variance of samples was not met, the

Wilcoxon matched-pairs signed rank test or Mann-Whitney U test was performed. More detailed statistical information is provided in Table S1.

## Materials Availability

This study did not generate new unique reagents.

## Data and code availability

Electrophysiological recording data have been deposited at ZENODO [data-type-specific repository] as [Database: accession number] and are publicly available as of the date of publication. All original code has been deposited at Github and is publicly available as of the date of publication. Any additional information required to reanalyze the data reported in this paper is available from the lead contact upon request.

## Acknowledgments

This work was funded by the European Research Council (ERC) under the European Union’s Horizon 2020 research and innovation programme (grant agreement No 101003187) and by the “la Caixa” Foundation (ID 100010434), under the agreement LCF/PR/HR20/52400020. The work was also funded by a Bial Foundation grant (175/2020). Part of the work received funding from the FCT under the scope of the projects PTDC/MED-NEU/4804/2020 (https://doi.org/10.54499/PTDC/MED-NEU/4804/2020), PTDC/SAU-TOX/6802/2020 (https://doi.org/10.54499/PTDC/SAU-TOX/6802/2020), 2022.02201.PTDC (https://doi.org/10.54499/2022.02201.PTDC) and 2022.01467.PTDC (https://doi.org/10.54499/2022.01467.PTDC). BC and LP have Scientific Employment Stimulus contracts from the Portuguese Foundation for Science and Technology (FCT) and 2023.08896.CEECIND; CEECIND/03898/2020 (https://doi.org/10.54499/2020.03898.CEECIND/CP1600/CT0015); CEECINST/00077/2018 (https://doi.org/10.54499/CEECINST/00077/2018/CP1640/CT0003). Host laboratory is funded by National funds, through FCT - project UIDB/50026/2020 (https://doi.org/10.54499/UIDB/50026/2020), UIDP/50026/2020 (https://doi.org/10.54499/UIDP/50026/2020) and LA/P/0050/2020 (https://doi.org/10.54499/LA/P/0050/2020). This work was also supported by Centre National de la Recherche Scientifique (CNRS); Fédération pour la Recherche sur le Cerveau, and Rotary - Espoir en Tête (to JB) and FRM EQU202403018036 (JB); Institut National du Cancer (JB). LR is a recipient of a Ph.D. fellowship from the FRM. Additional support by the French government through the France 2030 investment plan managed by the National Research Agency (ANR), as part of the Initiative of Excellence of Université Côte d’Azur under reference number ANR-15-IDEX-01, and The Collegium of Advanced Studies to JB.

Authors would like to acknowledge Luke Sjulson and Eliezyer Fermino de Oliveira for the valuable discussions regarding silicon probes implantation and recordings.

## Legends to Supplementary Figures

**Figure S1.**
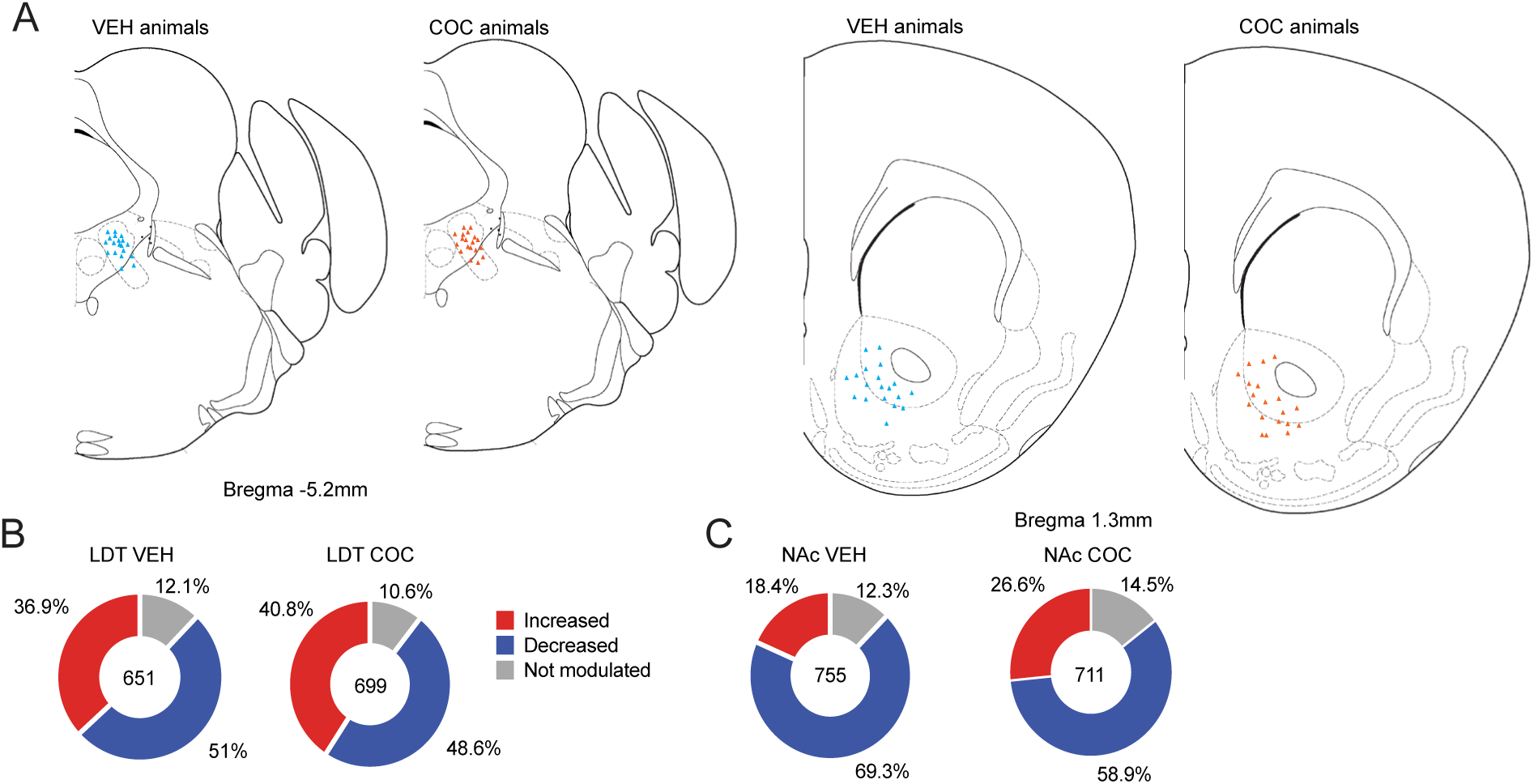
Histological probe placement and Percentages of responses in total LDT and NAc cells. **A)** Probe placement confirmation in VEH and COC animals in the LDT and NAc regions. Percentage of responses of LDT **(B)** and NAc **(C)** cells that increase, decrease or do not change their firing rate upon an acute dose of cocaine (10mg/Kg).

**Figure S2.**
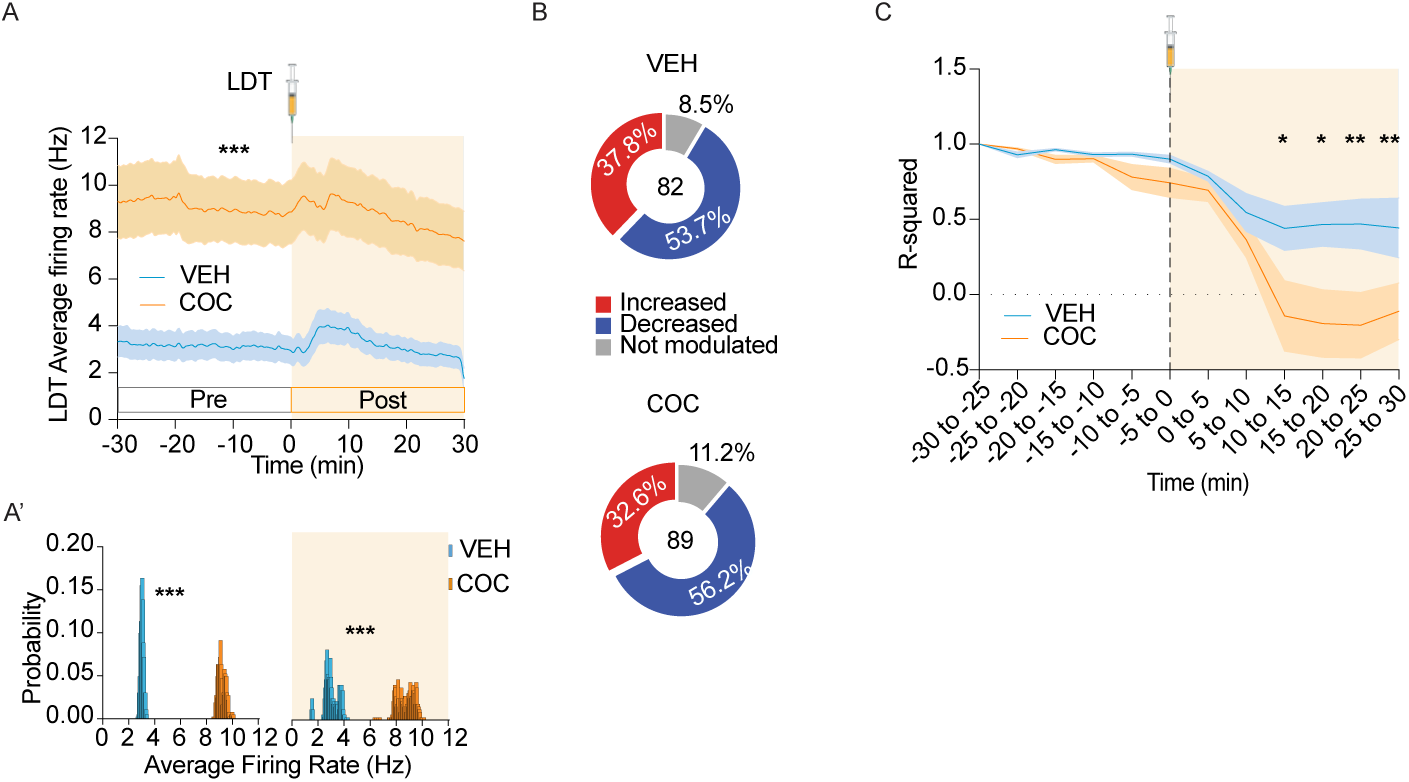
Neuroadaptations in the LDT are observed 15 days after last cocaine exposure. **A)** PSTH depicting the average firing rate of the LDT for the COC (orange, n=89) and VEH (blue, n=82) groups. Cocaine was administered at 0 timepoint. **A’** Distribution of the average firing rate in VEH and COC groups in the pre and post cocaine periods. Pre-exposure to cocaine increases average basal firing rate of the LDT. **B)** Percentage of responses of LDT cells that increase, decrease or do not change their firing rate upon an acute dose of cocaine (10mg/Kg). **C)** Baseline-trained RRR model (fitted to the first 5 min of pre-injection data) to sliding 5 min non-overlapping windows covering 30 min before and 30 min after injection. COC group decreases prediction upon cocaine exposure. Data represented as mean±SEM. *p < 0.05, **p < 0.01, ***p < 0.001.

**Figure S3.**
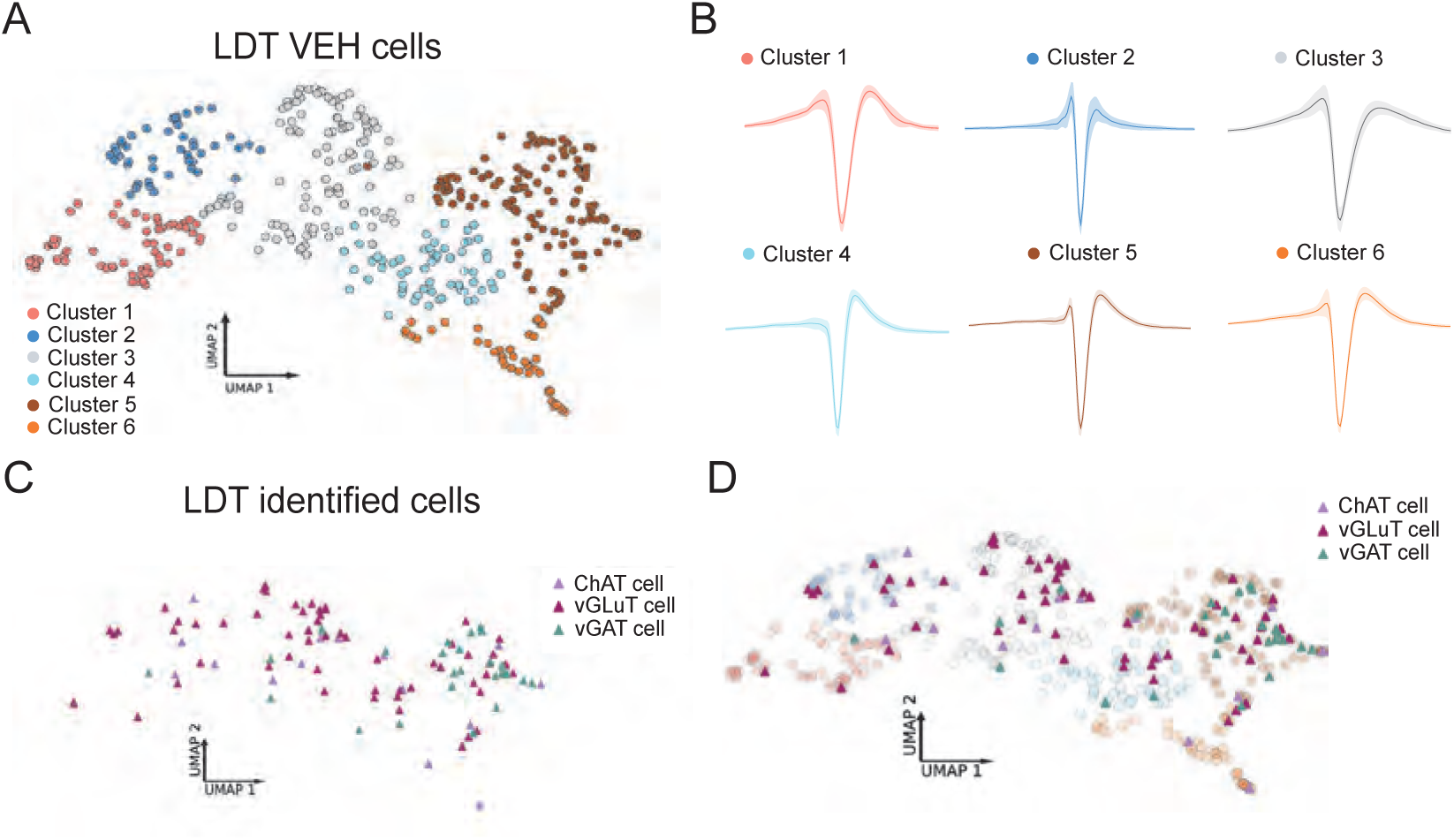
Clusterization of LDT neurons based on waveform characteristics. **A)** Analysis of waveform of LDT neurons from VEH animals using WaveMAP tool identified 6 neuronal clusters. Different colors represent distinct clusters based on waveform characteristics. **B)** Mean waveforms (solid lines) and standard deviations (shaded areas) for each of the six clusters. **C)** Optogenetically identified cholinergic (ChAT), glutamatergic (vGLUT2), and GABAergic (vGAT) LDT neurons. **D)** These distinct neuronal subtypes were distributed throughout more than one cluster, indicating diversity within each neuronal population.

**Figure S4.**
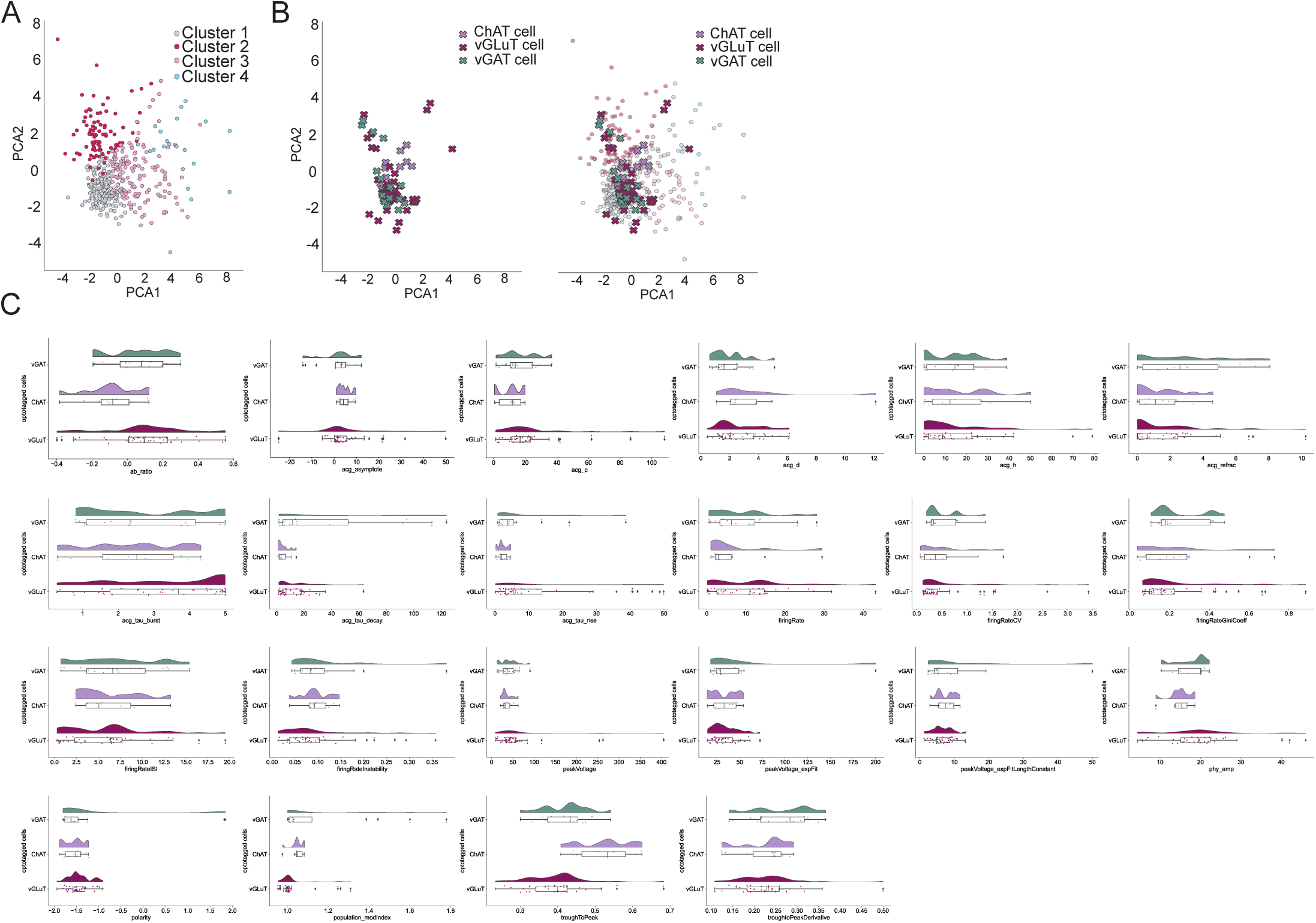
Clusterization of LDT neurons based on electrophysiological parameters. **A)** Using CellExplorer tool, we extracted 22 electrophysiological parameters of recorded LDT neurons. We then applied Principal Component Analysis (PCA) followed by clustering using the K-means algorithm. This approach identified four distinct neuronal clusters. **B)** Cholinergic, glutamatergic and GABAergic optotagged cells were distributed throughout more than one cluster, indicating diversity within each neuronal population. **C)** We performed Raincloud plot analysis using diverse combinations of CellExplorer parameters in order to identify those that could be used to segregate neuronal populations. GABAergic, glutamatergic and cholinergic cells’ distribution was overlapping for most of the parameters.

**Figure S5.**
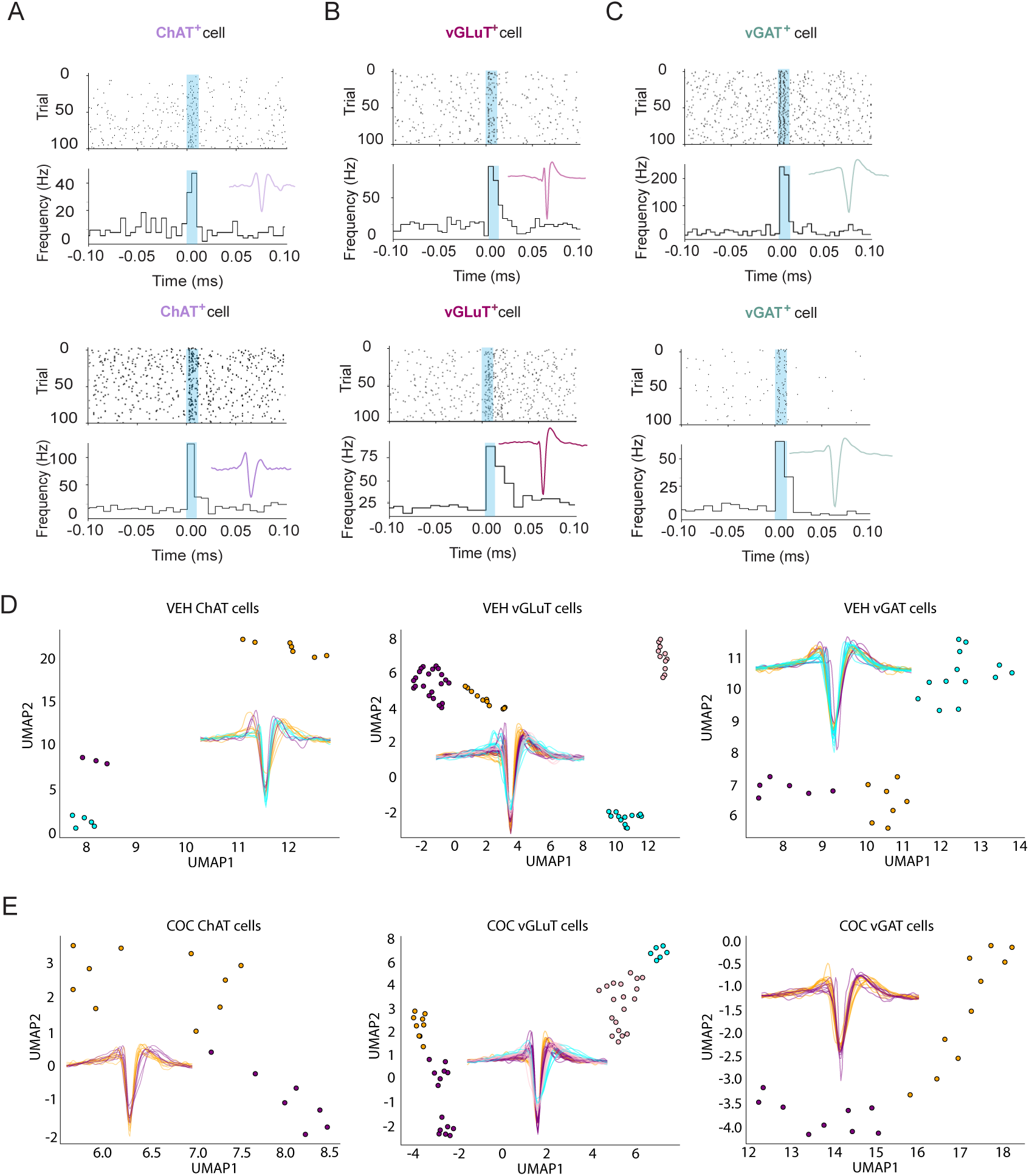
Subclusters within cholinergic, GABAergic and glutamatergic neuronal populations. Example of genetically-identified **A)** ChAT, **B)** vGLuT2 and **C)** vGAT cells in the LDT. As it can be observed in all subtypes, top cells are clearly distinct from bottom cells, though they are “genetically-identical” for the abovementioned markers. **D)** In line, we were able to identify several subclusters within each neuronal type, using WaveMap toolset. VEH ChAT neurons were divided into 3 clusters; vGLuT neurons grouped into 3 subclusters, and 4 subclusters of Vgat neurons. **E)** Remarkably, pre-exposure to cocaine alters the composition of these clusters. In COC animals, we found 2 ChAT, 2 vGluT2 and 4 vGAT subclusters.

**Figure S6.**
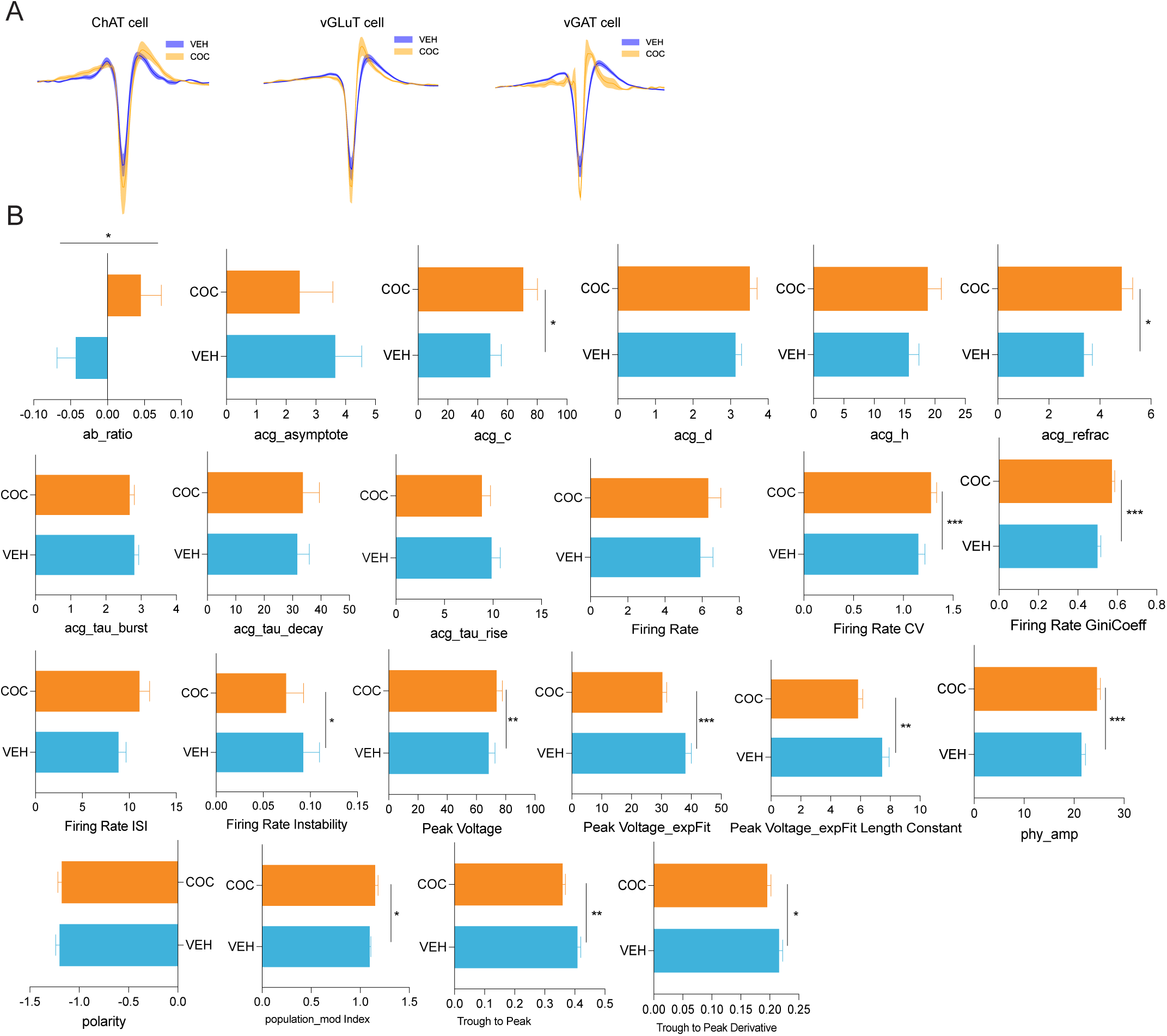
Cocaine pre-exposure triggers long-lasting changes in cholinergic, GABAergic and glutamatergic neuronal populations’ electrophysiological properties. **A)** Mean waveforms (solid lines) and s.e.m. (shaded areas) of cholinergic, glutamatergic and GABAergic neurons showing clear differences between VEH and COC groups, suggesting that pre-exposure to cocaine triggers long lasting changes in all populations. **B)** CellExplorer parameter analysis showed that pre-exposure to cocaine alters the electrophysiological properties of LDT neurons, limiting the direct comparison of VEH and COC clusters. Significant statistical differences were identified in the following parameters: ab_ratio, acg_c, acg_refrac, firing rate CV, firing rate GiniCoeff, firing rate instability, peak voltage, peak voltage_expFit, peak voltage_expFit Lenth Constant, phy_amp, population_mod Index, trough-to-peak, and trough-to-peak derivative. Data represented as mean±SEM. *p < 0.05, **p < 0.01, ***p < 0.001.

**Figure S7.**
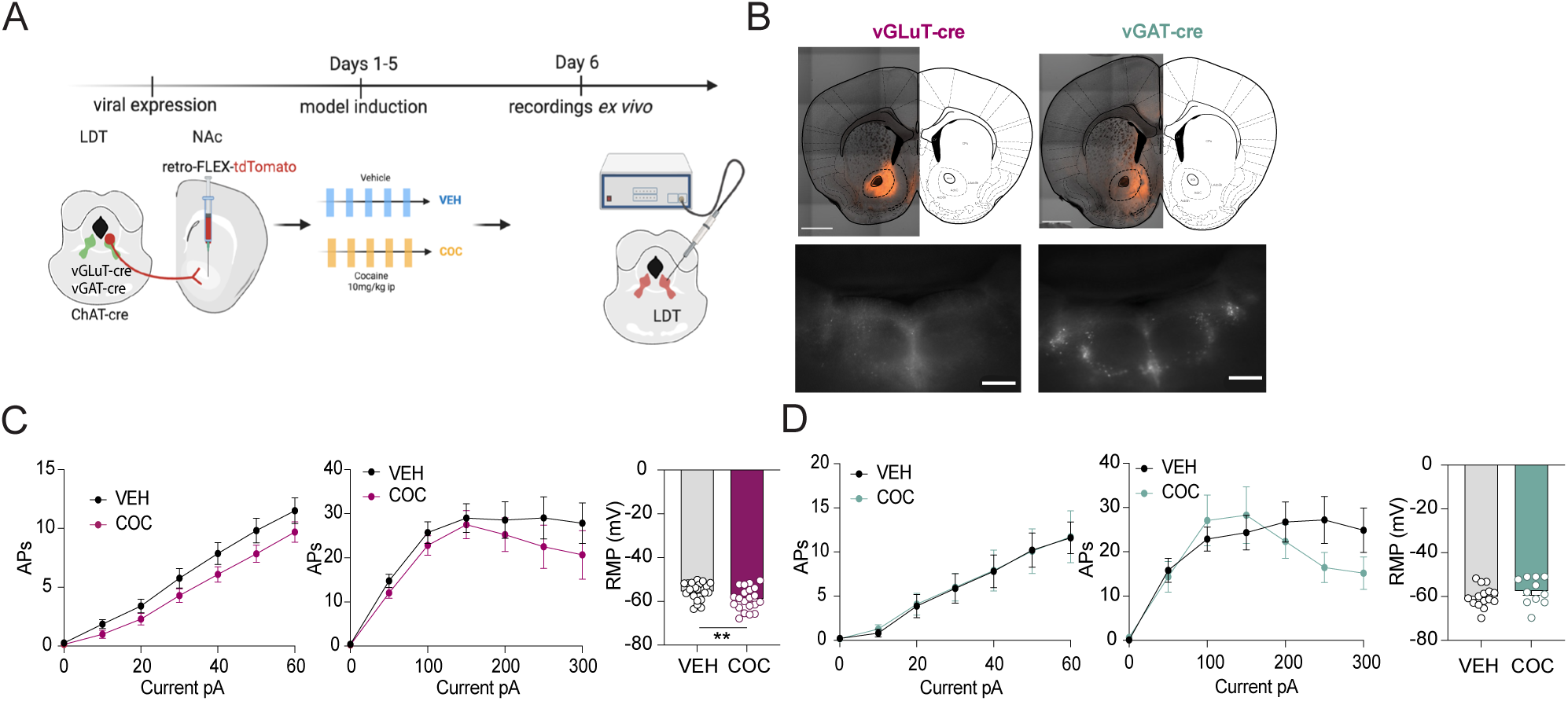
Cocaine pre-exposure does not robustly change LDT glutamatergic or GABAergic cells. **A)** Schematic experimental time line to perform patch-clamp recordings of LDT-NAc neurons. **B)** Example of retrograde virus injection in the NAc in vGluT2 and vGAT mice and visualization of tdtomato-tagged Glu and GABA neurons in the electrophysiological set-up during patch-clamp recordings. **C)** Cocaine exposure did not affect excitability or RMP in LDT glutamatergic (number of cell/mice: saline n = 22/5 and cocaine n = 20/5) or **D)** GABAergic cells (number of cell/mice: saline n = 15/3 and cocaine n = 12/3). Data represented as mean±SEM. *p < 0.05, **p < 0.01, ***p < 0.001.

**Figure S8.**
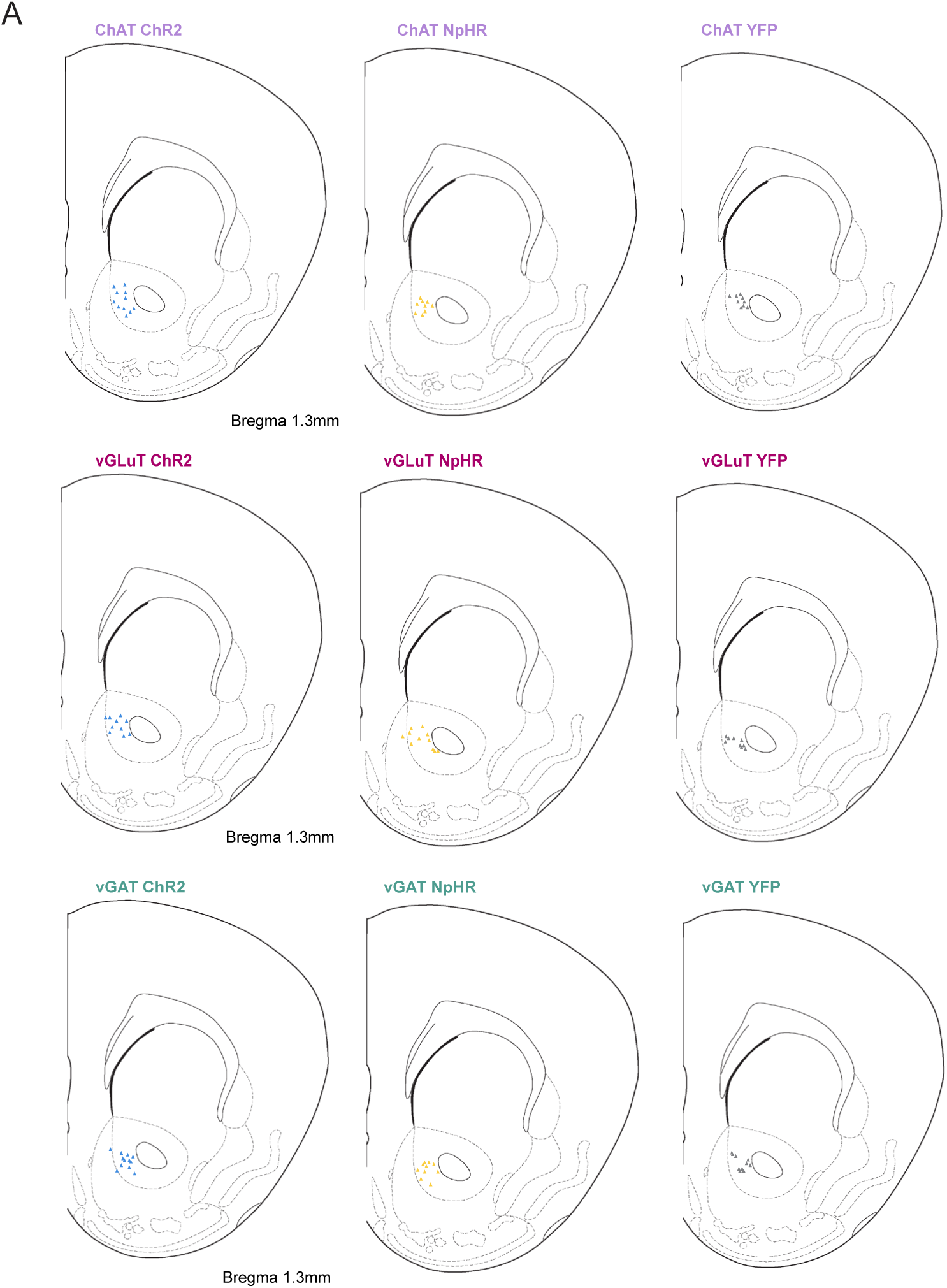
Fiber placement of ChAT-, vGLuT- and vGAT-cre animals. **A)** Location of fiber implant for ChAT-, vGLuT- and vGAT-ChR2; ChAT-, vGLuT- and vGAT-NpHR and ChAT-, vGLuT- and vGAT-YFP animals.

**Figure S9.**
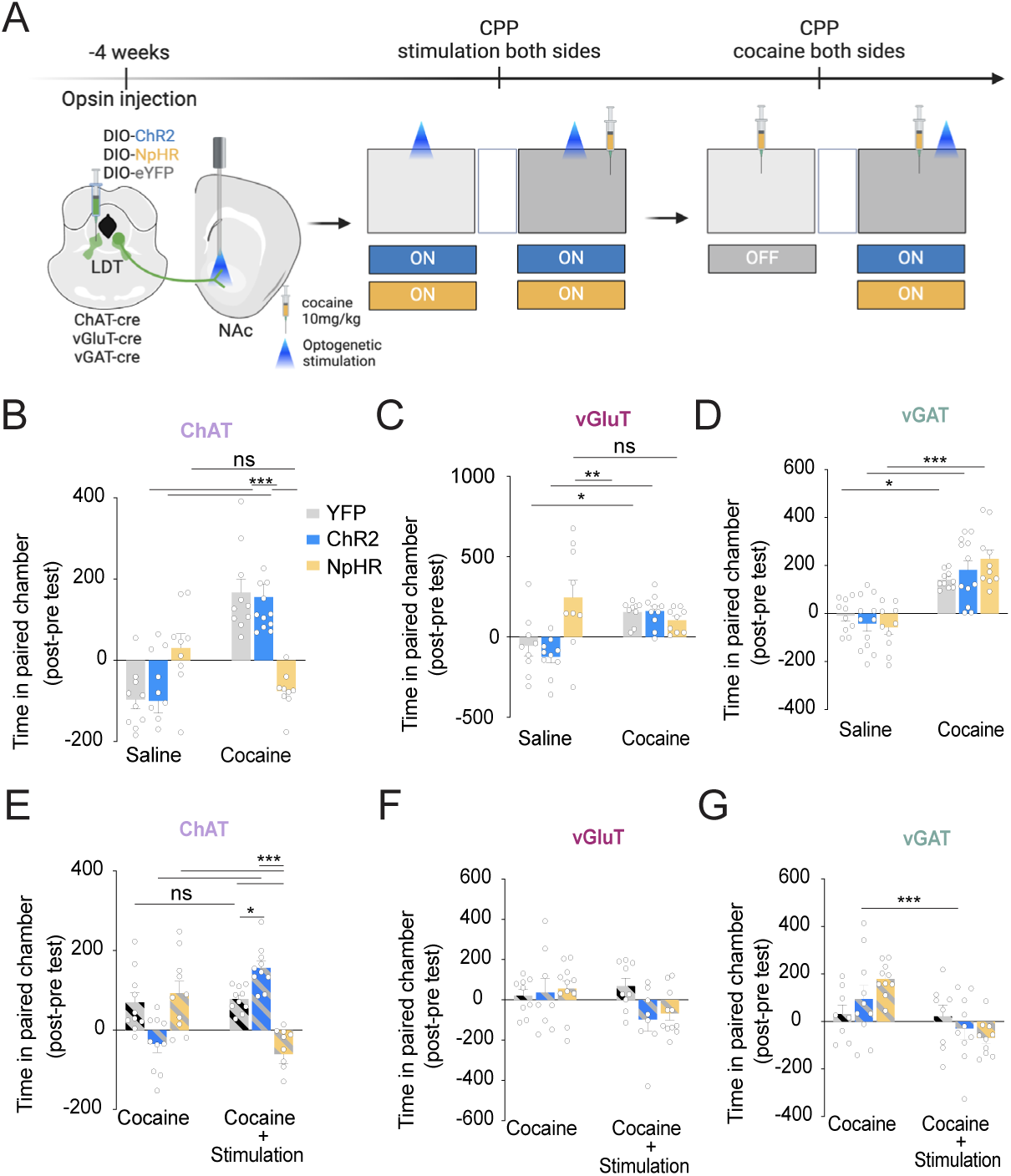
Modulation of LDT-NAc glutamatergic or GABAergic inputs does not impact cocaine conditioning. **A)** Schematic representation of the optogenetic manipulation experiments. ChAT-cre, vGluT2-cre or vGAT-cre animals were injected in the LDT with a cre-dependent ChR2 for excitation, or NpHR for inhibition or YFP as control. An optic fiber was placed in the NAc to allow stimulation of LDT terminals in this region. After opsin expression, optogenetic stimulation was performed in both sides of the apparatus and animals were conditioned to cocaine (10 mg/kg) to one side of the apparatus; saline was given to the other side (data in **B-D**). In another experiment, cocaine was given in the two sides of the apparatus to make them equally preferred to the animal, while optical stimulation was only applied to one of the sides (data in **E-G**). **B)** In the CPP with stimulation in both sides, optical activation of LDT-NAc cholinergic terminals had no effect on the observed preference for the cocaine-paired chamber, as ChAT-ChR2 animals were similar to YFP animals in terms of preference score (n_ChAT-ChR2_=11; n_ChAT-NpHR_=9, n_ChAT-YFP_=10). However, inhibition of LDT-NAc cholinergic neurons significantly reduced the preference for the cocaine-paired chamber. **C)** No effect in time spent in cocaine chamber was found between groups when LDT-NAc glutamatergic inputs were activated or inhibited (n_glut-ChR2_=10; n_glut-NpHR_=9, n_glut-YFP_=9). **D)** Modulation of LDT-NAc GABAergic inputs did not impact cocaine place preference (n_GABA-ChR2_=12; n_BAGA-NpHR_=10, n_GABA-YFP_=10). **E)** In the CPP with cocaine in both sides and stimulation in just one of the sides, optical activation of LDT-NAc cholinergic terminals resulted in significantly more time spent in the stimulated chamber compared to the non-stimulated side. In contrast, inhibition of LDT cholinergic neurons during cocaine conditioning significantly decreased time spent in the stimulated chamber (n_ChAT-ChR2_=10; n_ChAT-NpHR_=10, n_ChAT-YFP_=10). **F)** Optical excitation or inhibition of glutamatergic LDT-NAc inputs did not affect preference for either chamber (n_glut-ChR2_=8; n_glut-NpHR_=11, n_glut-YFP_=9). **G)** Optical excitation of GABAergic LDT-NAc inputs did not affect preference for either chamber. Inhibition of GABAergic LDT-NAc inputs slightly decreased preference for the stimulated side (n_GABA-ChR2_=10; n_BAGA-NpHR_=10, n_GABA-YFP_=8). Data represented as mean±SEM. *p < 0.05, **p < 0.01, ***p < 0.001.

